# Molecular mechanism of phospholipid transport at the bacterial outer membrane interface

**DOI:** 10.1101/2022.06.08.495403

**Authors:** Jiang Yeow, Min Luo, Shu-Sin Chng

## Abstract

The outer membrane (OM) of Gram-negative bacteria is an asymmetric lipid bilayer with outer leaflet lipopolysaccharides (LPS) exposed to extracellular milieu and inner leaflet phospholipids (PLs) facing the periplasm. This unique lipid asymmetry is the key to its innate drug resistance, rendering the OM impermeable to external insults, including antibiotics and bile salts. To maintain this OM barrier, the OmpC-Mla system removes mislocalized PLs from the OM outer leaflet, and transports them back to the inner membrane (IM); in the first step, the OM OmpC-MlaA complex transfers PLs to the periplasmic chaperone MlaC. This process likely occurs via a hydrophilic channel in MlaA, yet mechanistic details have remained elusive. Here, we biochemically and structurally characterize the architecture of the MlaA-MlaC transient complex. We map the interaction surfaces between MlaA and MlaC in *Escherichia coli*, revealing that MlaC binds MlaA at the periplasmic face in a manner that possibly juxtaposes the MlaA channel and the MlaC lipid binding cavity. In addition, we show that electrostatic interactions between the putative C-terminal tail helix of MlaA and a surface patch on MlaC are important for recruitment of the latter to the OM. We further provide biochemical evidence for conformational changes in the MlaA channel that correlate with interactions with MlaC and OM porins, as well as functional states of MlaA. Finally, we solve a 2.9-Å cryo-EM structure of OmpC-MlaA in nanodiscs in a disulfide-trapped complex with MlaC, reinforcing the mechanism of MlaC recruitment, and highlighting membrane thinning as a plausible strategy for directing lipids into the MlaA channel. Our work offers critical insights into how the OmpC-MlaA complex catalyzes retrograde transport of PLs to the IM to maintain OM lipid asymmetry.

## Introduction

Gram-negative bacteria assemble two lipid bilayers in their cell envelopes - the inner membrane (IM) and the outer membrane (OM) that encapsulate the cytoplasm and periplasm, respectively. At the OM, lipopolysaccharides (LPS) pack tightly with divalent cations and occupy the outer leaflet, while phospholipids (PL) reside mostly in the inner leaflet, presenting a unique bilayer with extreme asymmetry (Funahara & Nikaido, 1980; Kamio & Nikaido, 1976; Nikaido, 2003). The outer leaflet of LPS forms a stable layer with drastically reduced permeability, offering an effective protective barrier against the entry of toxic compounds, such as detergents and antibiotics. Proper establishment and maintenance of lipid asymmetry is thus critical for OM barrier function, which allows Gram-negative bacteria to survive in the most hostile environments, including the mammalian intestinal tract.

How OM lipid asymmetry is achieved has been extensively studied. It is well known that LPS are transported and inserted into the outer leaflet of the OM via the Lpt protein bridge machinery (Okuda, Sherman, Silhavy, Ruiz, & Kahne, 2016). Other major OM components, i.e. β-barrel proteins and OM lipoproteins, are assembled into the membrane via the Bam (Konovalova, Kahne, & Silhavy, 2017) and Lol (Okuda & Tokuda, 2011) pathways, respectively. It is less understood how exactly PLs are brought to the OM (Yeow & Chng, 2022), but recent studies suggest the involvement of a redundant set of AsmA-like proteins in this process (Douglass, McLean, & Trent, 2022; Grimm et al., 2020; Ruiz, Davis, & Kumar, 2021). Evidently, balancing the levels of all these OM components during active cell growth and division would be critical to ensure OM stability, and hence lipid asymmetry. In this regard, the Tol-Pal complex has been implicated in lipid homeostasis, possibly via removal of excess PLs from the OM (Shrivastava, Jiang, & Chng, 2017). When coordination across these pathways is sub-optimal, PLs may become mislocalized to the outer leaflet of the OM, forming PL bilayer patches that compromise OM lipid asymmetry and barrier function (Jia et al., 2004; Wu et al., 2005). To cope with these defects, additional cellular mechanisms are deployed to eliminate PLs that end up aberrantly in the outer leaflet of the OM. The OM phospholipase, PldA, hydrolyzes acyl chains from these PLs (Dekker, 2000), while the OM acyltransferase, PagP, transfers an acyl chain from outer leaflet PLs to LPS and phosphatidylglycerol (PG) (Bishop, 2005; Dalebroux, Matamouros, Whittington, Bishop, & Miller, 2014). Finally, the OmpC-Mla system, an established PL trafficking pathway, extracts PLs from the outer leaflet of the OM and shuttles them back to the IM (Chong, Woo, & Chng, 2015; Malinverni & Silhavy, 2009).

The OmpC-Mla system, first identified in *Escherichia coli* based on homology to the TGD2 pathway in *Arabidopsis thaliana*, comprises the OM lipoprotein MlaA in complex with osmoporin OmpC, the periplasmic chaperone MlaC, and the IM ATP-binding cassette (ABC) transporter MlaFEDB (Chong et al., 2015; Malinverni & Silhavy, 2009; Thong et al., 2016). Removing *ompC* or any *mla* component results in PL accumulation in the outer leaflet of the OM of *E. coli*, indicating a role in maintaining OM lipid asymmetry (Chong et al., 2015; Malinverni & Silhavy, 2009). To achieve that, the OmpC-MlaA complex somehow extracts PLs from the outer leaflet of the OM, hands them over to MlaC, which in turn transfers these PLs into the IM via the MlaFEDB complex (Low & Chng, 2021; Low, Thong, & Chng, 2021; Tang et al., 2021). Many recent biochemical and structural studies have provided detailed insights into ATP-dependent PL transfer steps at the IM (Chi et al., 2020; Coudray et al., 2020; Ekiert et al., 2017; Ercan, Low, Liu, & Chng, 2019; Hughes et al., 2019; Mann et al., 2021; Tang et al., 2021; Thong et al., 2016; Zhang, Fan, Chi, Zhou, & Li, 2020; Zhou et al., 2021). In particular, we now know that when PL-bound MlaC arrives at the IM, it can spontaneously transfer the lipid molecule to the binding cavity of MlaFEDB (Low et al., 2021). Even though this initial process is reversible (Hughes et al., 2019; Low et al., 2021; Tang et al., 2021), ATP binding/hydrolysis by the MlaFEDB complex ultimately catalyzes the transfer of PLs into the IM, driving overall retrograde transport from the OM (Low et al., 2021; Tang et al., 2021). How ATP binding/hydrolysis may be coupled to PL transport in the MlaFEDB complex can be partly inferred from recently solved structures (Chi et al., 2020; Coudray et al., 2020; Mann et al., 2021; Tang et al., 2021; Zhang et al., 2020; Zhou et al., 2021), though detailed mechanistic understanding is still limited.

Lipid transfer at the OM is somewhat less understood, especially how retrograde PL movement from the outer leaflet of the OM to MlaC can be facilitated in a manner not (directly) dependent on conventional energy sources at the IM. The OM lipoprotein MlaA interacts with trimeric porins OmpC and OmpF, where OmpC appears to play a more critical role in Mla function (Chong et al., 2015). It has been established that, unlike canonical lipoproteins, MlaA resides entirely within the OM bilayer and makes extensive contacts with porins at their subunit interfaces (Abellon-Ruiz et al., 2017; Yeow et al., 2018). Moreover, MlaA itself forms a hydrophilic channel in the OM; it comprises a ‘donut’-shaped architecture (constituted largely by five horizontal helices) and a vertical ‘ridge’ (formed by a ‘helix-turn-helix’ motif) defining the central channel. The MlaA channel may provide a plausible path for PL translocation across the OM, and may be gated by conformational movements of an adjacent β-hairpin loop motif. However, we do not yet have clear ideas of how MlaA actually sits in the OM lipid bilayer, as well as the expected conformational changes that should occur in the protein during PL transport. MlaC binds PLs with presumed high affinity (Ercan et al., 2019; Huang et al., 2016; Hughes et al., 2019), and has been demonstrated to interact with MlaA in vivo (Ercan et al., 2019) and in vitro (Ekiert et al., 2017). Yet it remains elusive how exactly MlaA interacts with MlaC to allow spontaneous, productive transfer of PLs.

Here, we elucidate a molecular picture of the transient MlaA-MlaC complex during PL transfer at the OM. Using in vivo photo- and disulfide-crosslinking, we demonstrate that MlaC interacts with MlaA directly at its periplasmic base, in a manner that juxtaposes the lipid binding cavity of MlaC with the hydrophilic channel of MlaA. In addition, mutational analyses reveal that MlaA recruits MlaC to the OM via additional electrostatic interactions between a C-terminal tail helix of MlaA and a surface charged patch on MlaC. We further establish solvent-accessibility changes in a key channel residue in MlaA that are influenced by interactions with MlaC and porins (OmpC/F), as well as functionality of MlaA itself. Finally, we solve the cryo-EM structure of a disulfide-trapped OmpC-MlaA-MlaC complex in a lipid bilayer, elucidating how MlaA recruits MlaC, and how it deforms the membrane to facilitate transport. Collectively, our work provides novel mechanistic insights into retrograde PL transfer at the bacterial OM critical for maintaining lipid asymmetry.

## Results

### MlaC interacts with α-helices at the periplasmic opening of the hydrophilic channel of MlaA

To receive PLs from MlaA, MlaC would have to gain access to the hydrophilic MlaA channel from the periplasmic side, albeit transiently. To map the region(s) on MlaA that contacts MlaC, we replaced ~27 residues on the periplasmic face of *E. coli* MlaA separately with *para*-L-benzoylphenylalanine (*p*Bpa) (Chin, Martin, King, Wang, & Schultz, 2002), and examined which positions allow photoactivatable crosslinks with MlaC. We found five positions on MlaA that demonstrated strong photoactivatable crosslinks with MlaC when substituted with *p*Bpa (**Fig. 1A**). Another five positions exhibited weaker crosslinks to MlaC (**Fig. S1**). Interestingly, these residues are all located on the C-terminal helices (Q195-N226) of MlaA extending from the channel opening in the periplasmic face (**Fig. 1B**), suggesting that MlaC can be juxtaposed to receive lipids from MlaA at the base of the channel.

**Figure 1.**
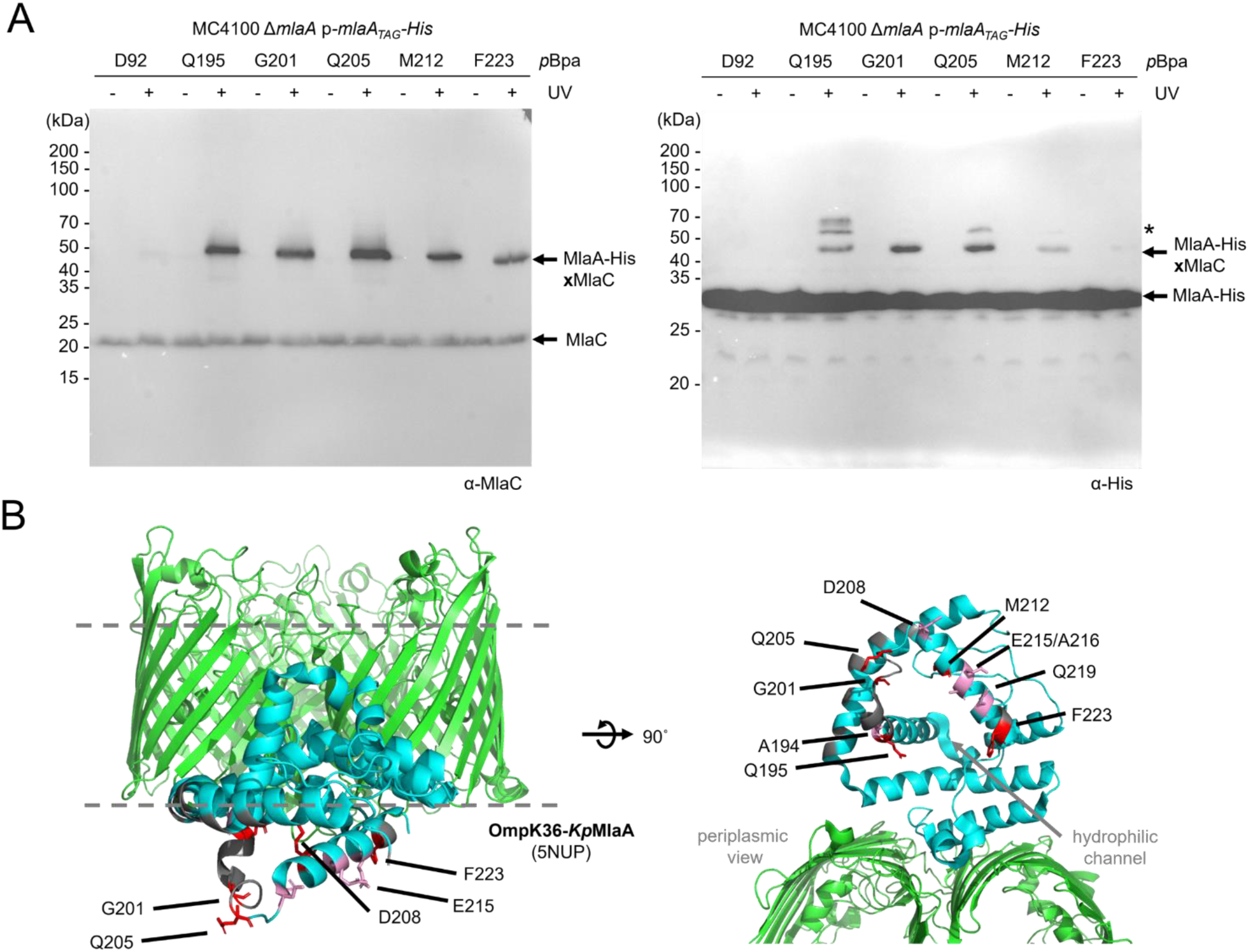
MlaA contacts MlaC in cells via C-terminal α-helices at the periplasmic opening of its hydrophilic channel. (A) Representative immunoblots showing UV-dependent formation of strong crosslinks between MlaA and MlaC in Δ*mlaA* cells expressing MlaA-His substituted with *p*Bpa at indicated positions from the pCDF plasmid. Additional but unidentified MlaA_*p*Bpa_-His crosslinked adducts were detected and denoted with an asterisk (*). (B) Membrane (*left*) and periplasmic (*right*) views of cartoon representations of the crystal structure of *Klebsiella pneumoniae* OmpK36 (*green*)-MlaA (*cyan*) (PDB 5NUP) (Abellon-Ruiz et al., 2017) with positions that crosslink to MlaC highlighted. Residues exhibiting strong, weak, or no photo-crosslinks with MlaC are illustrated in *red* and *pink* sticks, or in grey respectively. The OM boundaries are indicated as *gray* dashed lines.

We have also previously established the residues on MlaC that contact MlaA via the same photocrosslinking approach (Ercan et al., 2019). To determine reciprocal contact points between the two proteins, we introduced cysteines at positions on either protein that displayed strong photoactivatable crosslinks, and monitored disulfide bond formation between these MlaA and MlaC cysteine variants in cells. We tested 24 combinations of cysteine residue-pairs from four and six positions on MlaA and MlaC, respectively. Of these pairs, 11 enabled disulfide bond formation between MlaA and MlaC, indicating that these residue positions can lie in close proximity when the two proteins interact (**Fig. 2A**). MlaA^Q195C^ and MlaC^Q151C^ did not form disulfide crosslinks with any partner cysteine variants tested. In contrast, Q205, M212 and F223 on MlaA, and K128, S172 and T175 on MlaC, each allowed disulfide bond formation with multiple tested positions on their partner protein, when replaced with cysteine (**Figs. 2A and 2B**). The strongest disulfide crosslink was MlaA^Q205C^-MlaC^V171C^ (**Figs. 2A and S2**), enabled perhaps by sufficiently long lifetime(s) of a transient interaction mode in the complex. Interestingly, Q205 lies directly at the base of the hydrophilic channel of MlaA while V171 is positioned at the entrance of the lipid binding cavity of MlaC (**Fig. 2B**), suggesting that the trapped MlaA-MlaC complex may be close to or approaching an arrangement(s) that could eventually lead to productive lipid transfer.

**Figure 2.**
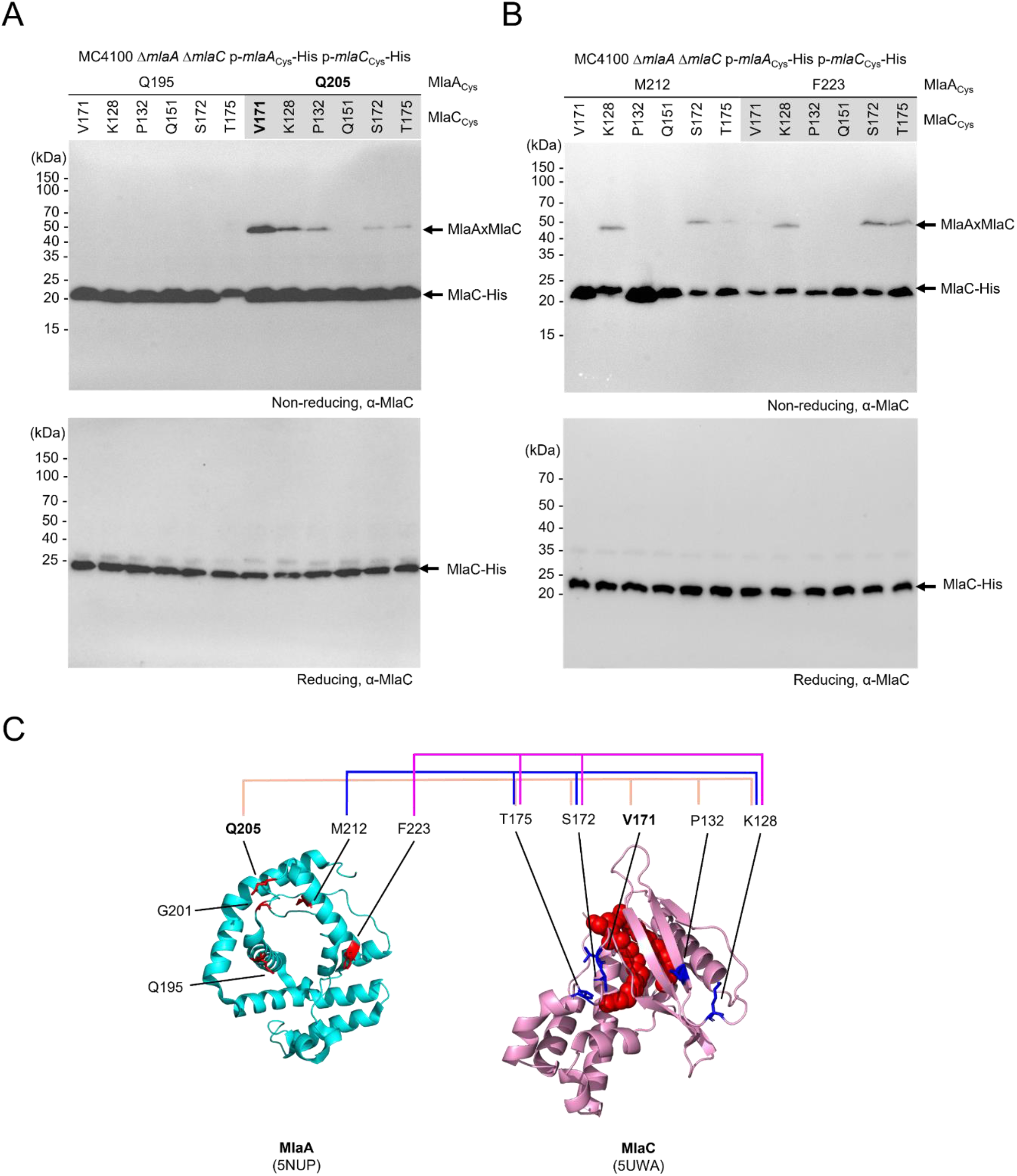
MlaC interacts with MlaA in a manner that juxtaposes the MlaA channel and the MlaC lipid binding cavity. (A) Representative immunoblots showing formation of disulfide crosslinks between MlaA and MlaC in Δ*mlaA* Δ*mlaC* cells expressing cysteine-substituted MlaA-His and MlaC-His from the pCDF and pET22/42 plasmids, respectively. Samples were subjected to non-reducing (*top*) or reducing (*bottom*) SDS-PAGE prior to immunoblotting. (C) Cartoon representation of *Kp*MlaA (*cyan*) (PDB 5NUP) (Abellon-Ruiz et al., 2017) and holo-MlaC (*pink*, PL in *red* spheres) (PDB 5UWA) (Ekiert et al., 2017). Residues on MlaC which displayed disulfide crosslinking with MlaA, and vice versa, are illustrated in *red* and *blue* sticks, respectively. MlaC residues mapped to MlaA residues Q205C, M212C and F223C are connected by *beige, blue* and *pink lines* respectively.

### Electrostatic interactions between the C-terminal tail helix of MlaA and a surface charged patch on MlaC are necessary for MlaC recruitment

We have shown that MlaC interacts with the C-terminal α-helices at the base of MlaA. However, there is presumably an unstructured region beyond these interacting helices (G227-E251; *E. coli* numbering) missing from the reported MlaA crystal structures (Abellon-Ruiz et al., 2017); yet, in the recent structural model predicted by AlphaFold (AF-P76506-F1) (Jumper et al., 2021), the extreme end (N238-E251) of this ‘tail’ region was confidently modelled as a short α-helix (**Fig. 3A**). To understand if this C-terminal tail helix is also important for MlaC interaction, we made several MlaA deletion constructs, and probed for MlaC interaction. While removing residues after P232 (MlaA^CTD4^) conferred milder SDS/EDTA sensitivity than cells lacking MlaA (**Fig. S3A**), it had completely abolished disulfide bond formation between MlaA^Q205C^ and MlaC^V171C^ (**Fig. 3B**). In fact, deletion of the last 9 amino acids on MlaA^D243X^ (MlaA^CTD6^), a highly negatively charged segment on the putative terminal helix, was sufficient to disrupt disulfide crosslinks with MlaC. To test the importance of electrostatics, we also constructed two sets of Arg mutants at a single face of this helix, MlaA^D243R/D244R^ (MlaA^2DD2R^) and MlaA^D247R/E251R^ (MlaA^2DE2R^) (**Fig. 3A**), and found that these charge-reversed variants drastically reduced disulfide crosslinks between MlaA^Q205C^ and MlaC^V171C^ (**Fig. 3B**). The different MlaA variants are still able to pull down trimeric OmpC, indicating they preserve overall MlaA structure (**Fig. S4**). We conclude that the negatively-charged residues on the C-terminal tail helix of MlaA are also critical for MlaC interaction.

**Figure 3.**
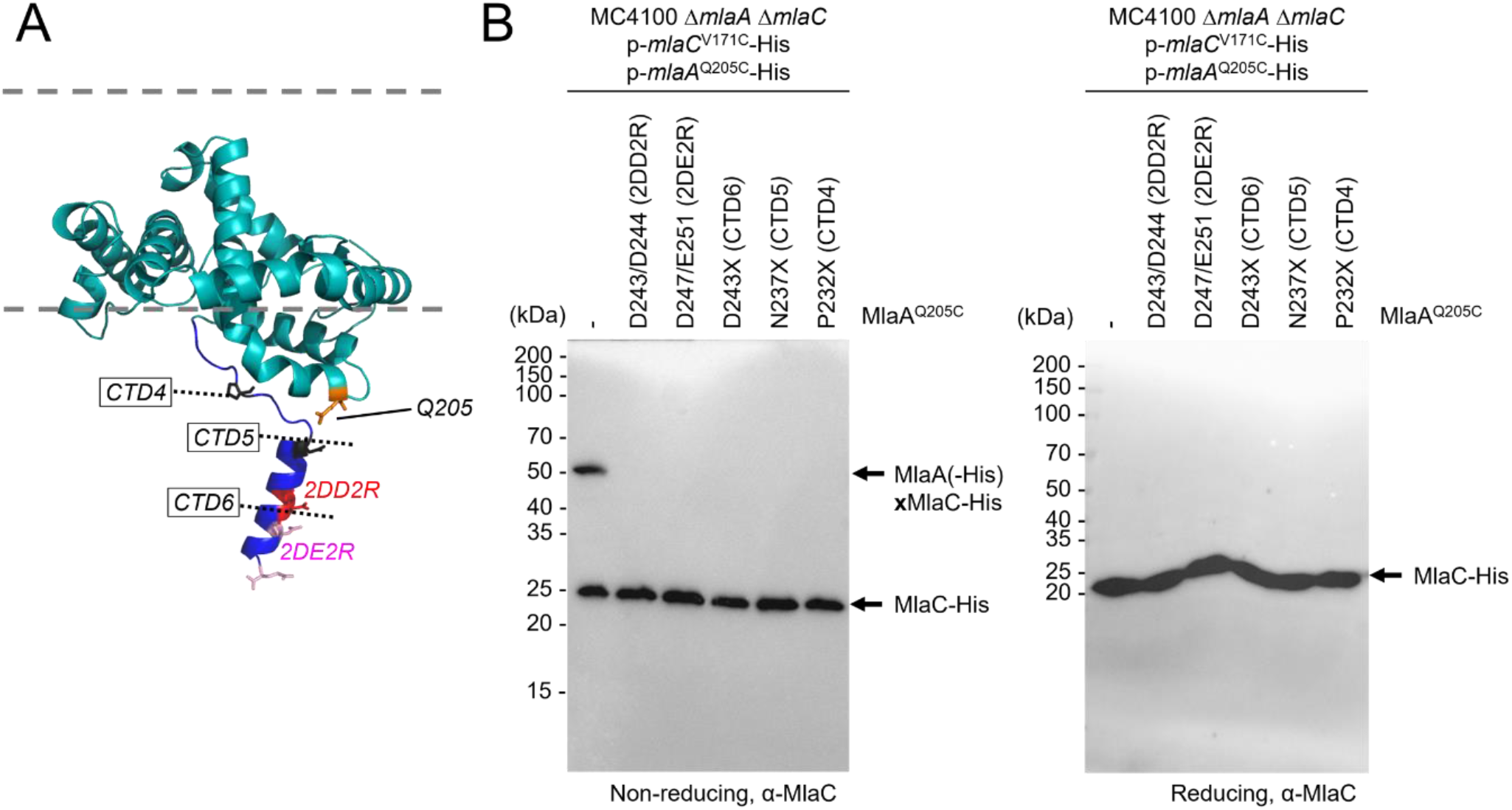
Negatively-charged residues on the putative C-terminal tail helix of MlaA are important for MlaC recruitment. (A) Cartoon representation of the *Ec*MlaA (*teal*) model generated from AlphaFold2 (AF-P76506-F1) (Jumper et al., 2021) with the putative C-terminal tail helix coloured *blue*. C-terminal truncation positions (CTDs) are annotated. MlaC-crosslinking residue Q205 is labelled in *orange* sticks. Charge-reversed residues in MlaA^2DD2R^ and MlaA^2DE2R^ are labelled in *red* and *pink* sticks respectively. (B) Representative immunoblots showing abolishment of disulfide crosslinks between MlaA and MlaC in Δ*mlaA* Δ*mlaC* cells expressing various C-terminal tail helix mutants of MlaA^Q205C^(-His) and MlaC^V171C^-His from the pCDF and pET22/42 plasmids, respectively. Samples were subjected to non-reducing (*left*) or reducing (*right*) SDS-PAGE prior to immunoblotting.

We next asked if there are specific positively-charged regions on MlaC that interact with the C-terminal tail helix of MlaA. We identified seven clusters of positively-charged residues (Arg/Lys) all around the structure of apo-MlaC (Hughes et al., 2019) (**Fig. 4A**), and successfully converted five of these clusters to aspartates separately. MlaC^K84D/R90D^ (MlaC1) abolished disulfide crosslinks between MlaA^Q205C^ and MlaC^V171C^, while MlaC^K71D^ (MlaC0) and MlaC^R143D/R147D^ (MlaC3) led to partial loss (**Fig. 4B**), confirming the overall importance of electrostatics. These mutations did not introduce any SDS/EDTA sensitivity (**Fig. S3B**), however, suggesting they may only impact recruitment efficiency, but not overall lipid transport function. Remarkably, combining MlaA^D243R/D244R^ (MlaA^2DD2R^) with MlaC^K84D/R90D^ (MlaC1) fully restored disulfide bond formation (**Fig. 4C**), implying that residues D243/D244 on MlaA directly contact residues K84/R90 on MlaC during MlaA-MlaC interaction.

**Figure 4.**
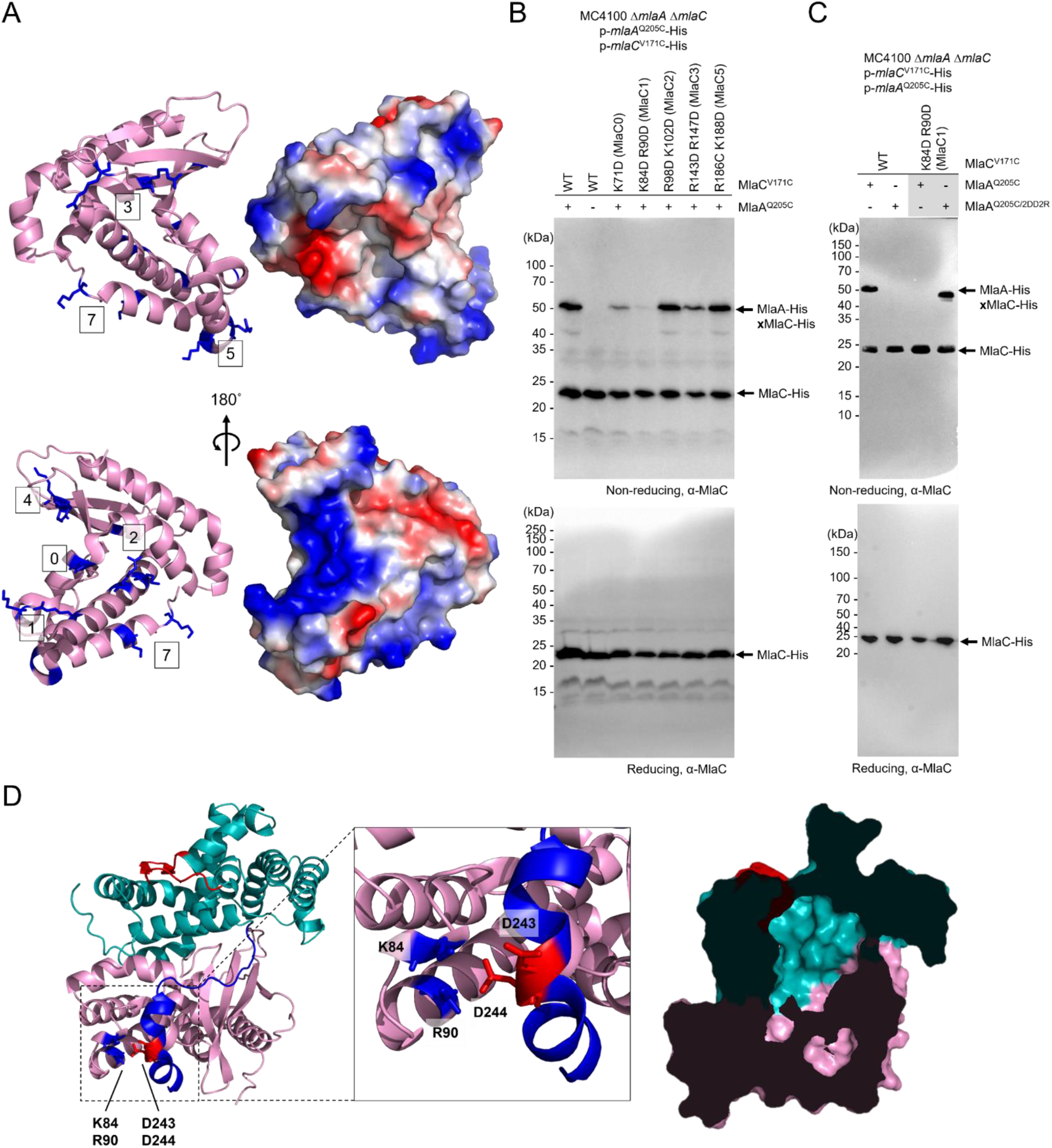
Positively-charged surface residues on MlaC are responsible for mediating MlaA interaction. (A) Cartoon and surface vacuum electrostatics representations of apo-MlaC (PDB 6GKI) (Hughes et al., 2019) revealing positively-charged patches (*blue surface*). Positively-charged residues (or groups of residues, *blue sticks*) are numbered from 0 to 7. (B) Representative immunoblots showing abolishment of disulfide crosslinks between MlaA and MlaC in Δ*mlaA* Δ*mlaC* cells expressing indicated surface charged-reversed mutants plasmids, respectively. (C) Representative immunoblots showing restoration of disulfide crosslinks between MlaA and MlaC in Δ*mlaA* Δ*mlaC* cells expressing charge-reversed MlaC1-His and MlaA^2DD2R^-His from pET22/42 and pCDF plasmids, respectively. Samples were subjected to non-reducing (*top*) or reducing (*bottom*) SDS-PAGE prior to immunoblotting. (D) Cartoon representations of Alphafold2-predicted MlaA (*teal*)-MlaC (*pink*) model, revealing the proximity of residues D243/D244 on MlaA, and residues K84/R90 on MlaC illustrated in *blue* and *red* sticks on MlaA and MlaC respectively (inset). Cutaway surface representation of this MlaA-MlaC model is shown on the right, revealing the juxtaposition of the hydrophobic lipid-binding cavity of MlaC with the hydrophilic channel of MlaA.

To obtain a possible picture of the MlaA-MlaC interface, we applied AlphaFold2 to predict the structural model of the MlaA-MlaC complex (Evans et al., 2021; Jumper et al., 2021; Mirdita et al., 2022). Amazingly, the model not only revealed an arrangement of MlaA and MlaC that is fully consistent with our in vivo *p*Bpa crosslinking data, it also correctly predicted the charge-charge interactions between the D243/D244 on MlaA and K84/R90 on MlaC (**Fig. 4D**). The C-terminal tail helix of MlaA is connected to the rest of the protein via a flexible linker, suggesting that this tail helix might serve as a bait to recruit apo-MlaC to the OM complex via electrostatic attraction. Importantly, once MlaC is stably bound and positioned, the hydrophilic channel in MlaA becomes continuous with the lipid binding cavity of MlaC (**Fig. 4D**), providing a glimpse into the route for PL transfer from the outer leaflet of the OM to MlaC.

### Interactions with MlaC modulate conformational changes in the MlaA channel

Productive MlaA-MlaC interactions would be required for PL transfer from the OmpC-MlaA complex to MlaC. We hypothesize that such interactions could elicit conformational changes in MlaA to facilitate PL transfer. To test this idea, we examined the solvent accessibilities, as an indication of possible conformational states, of specific residues within the MlaA channel in the absence of MlaC (**Fig. 5A**). We have previously shown using the substituted cysteine accessibility method (SCAM) that many hydrophilic residues in the MlaA channel are fully accessible to a membrane-impermeable thiol-reactive reagent ((2-sulfonatoethyl) methanethiosulfonate; MTSES) in wild-type cells (Yeow et al., 2018). This is true for additional channel residues identified as part of a more systematic scan for solvent accessibility (**Fig. S5**). One exception is K184, which exhibited partial MTSES accessibility when substituted with cysteine; within the MlaA population in wild-type cells, K184C is accessible to MTSES in some MlaA molecules but not others, in comparable proportions (**Fig. 5B**). Interestingly, removing MlaC resulted in K184C becoming MTSES-inaccessible in the entire population of MlaA (**Fig. 5B**), an effect not seen for other channel residues tested (**Fig. S5**). This shift in accessibility of K184C was reversed by expressing wild-type MlaC in trans, but not the charge-reversed MlaC1 variant that disrupted MlaA-MlaC interactions (**Fig. 5C**). This suggests that interaction with MlaC is required for MlaA to adopt the state where K184C is solvent accessible. Furthermore, we observed that K184C in MlaA also became MTSES-inaccessible in the Δ*mlaD* strain (**Fig. 5B**). While MlaC is still present in this strain, it is likely bound to PLs (i.e. holo form), given that off-loading of MlaC is not possible in the absence of an intact ABC transporter in the IM. Overall, our results indicate that the binding and release of apo-MlaC may influence the solvent accessibilities and conformational states of the MlaA channel.

**Figure 5.**
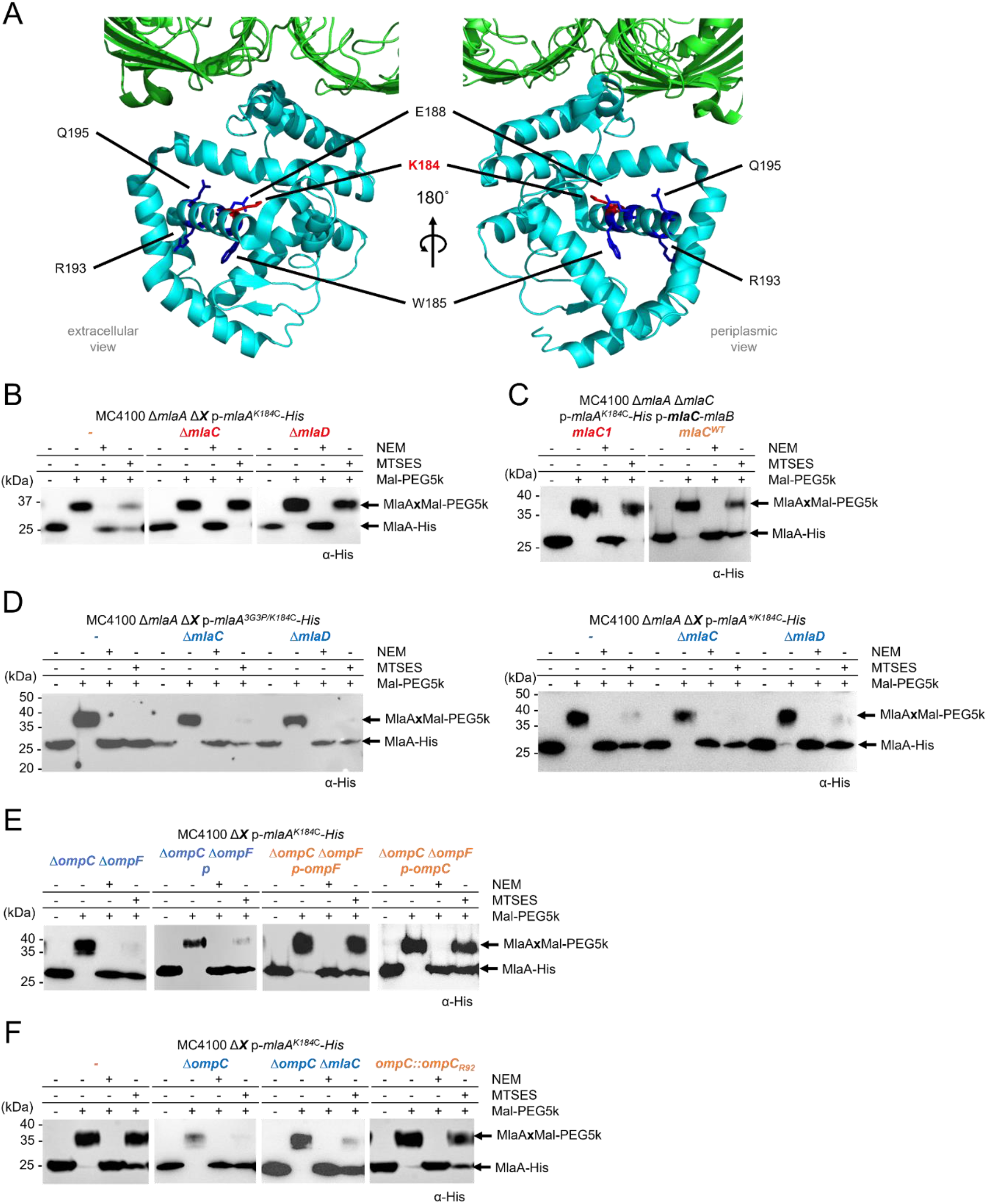
The MlaA hydrophilic channel exhibits solvent-accessibility changes due to interactions with MlaC and trimeric porins. Extracellular (*left*) and periplasmic (*right*) views of cartoon representations of the reported crystal structure of OmpK36 (*green*)-*Kp*MlaA (*cyan*) (PDB 5NUP) (Abellon-Ruiz et al., 2017), showing channel residues substituted with cysteine (for SCAM) in *sticks*. (B, C) Representative immunoblots showing Mal-PEG5k alkylation of MlaA^K184C^-His cysteine variant expressed from the pCDF plasmid, either in (B) Δ*mlaA*, Δ*mlaA* Δ*mlaC*, and Δ*mlaA* Δ*mlaD* backgrounds, or (C) Δ*mlaA* Δ*mlaC* background also producing wildtype or charge-reversed K84D/R90D (MlaC1) MlaC variants from the pET23/43-*mlaCB* plasmid. (D) Representative immunoblots showing Mal-PEG5k alkylation of MlaA^K184C^-His variant also harboring non-functional 3G3P or gain-of-function *mlaA\** mutations expressed from pET23/42 plasmids in Δ*mlaA*, Δ*mlaA* Δ*mlaC*, and Δ*mlaA* Δ*mlaD* background strains. (E, F) Representative immunoblots showing Mal-PEG5k alkylation of MlaA^K184C^-His cysteine variant expressed from the pCDF plasmid in various porin mutant backgrounds. In (E), OmpC or OmpF were expressed from pDSW206 plasmids where indicated. In SCAM, cells were labelled with membrane-permeable (NEM) or impermeable (MTSES) reagents, followed by alkylation with Mal-PEG5k, which introduces a ~5-kDa mass shift to MlaA^K184C^-His. The levels of solvent accessibility of K184C in MlaA in the various strains, i.e. fully, partially, or not blocked by MTSES, are highlighted in *blue*, *orange*, or *red*, respectively.

We have also considered the possibility that shifts in MTSES-accessibility could be due to OM asymmetry defects in these *mla* strains. This is unlikely, however, since K184C MTSES-accessibility in other OM-defective strains (Δ*tolB*, Δ*pal*) (Shrivastava et al., 2017) was similar to that in wild-type cells (**Fig. S6A**). Supporting this observation, the overexpression of phospholipase PldA, which can rescue OM asymmetry defects in *mla* mutants (Malinverni & Silhavy, 2009), was unable to reverse the observed shifts in accessibility of K184C in both Δ*mlaC* and Δ*mlaD* strains (**Fig. S6B**). We conclude that accessibility changes in K184 in MlaA are not due to the general loss of lipid asymmetry in the OM.

### Non-functional MlaA variants adopt conformational states that no longer respond to interactions with MlaC

It is intriguing that K184C exhibits accessibility changes that may reflect different conformational states in the MlaA channel, possibly due to functional interactions with apo-MlaC. We next asked whether this same residue can also help us discern the functionality of MlaA itself. We first examined the MTSES-accessibility of K184C in known MlaA variants. MlaA^G141P/G143P/G145P^ (MlaA^3G3P^) is a non-functional variant believed to have restricted flexibility in a β-hairpin loop that may gate the MlaA hydrophilic channel (Yeow et al., 2018). MlaAΔN43F44 (MlaA*) is a gain-of-function variant that is thought to have poor control of loop dynamics, resulting in PL flipping from the inner to the outer leaflet of the OM (Grimm et al., 2020; Sutterlin et al., 2016; Yeow et al., 2018). Interestingly, K184C in the entire population of MlaA^3G3P^ or MlaA* appeared to be fully accessible to MTSES in cells, contrary to that observed in wild-type MlaA (**Fig. 5D**). Furthermore, removing MlaC or MlaD did not alter the accessibility of K184C in both MlaA variants, suggesting that MlaA^3G3P^ and MlaA* are in conformational states that cannot respond to interactions with apo-MlaC. Both MlaA^3G3P^ and MlaA* are therefore non-functional in facilitating retrograde PL transfer to MlaC.

An outstanding question in the OmpC-Mla system relates to why only OmpC, but not OmpF, appears to function together with MlaA. Under high osmolarity conditions, OmpC is produced at higher levels than OmpF (Chong et al., 2015). Therefore, to gain insights into the role of porins in the pathway without confounding effects due to expression levels, we went on to examine the solvent accessibility of MlaA K184C in the Δ*ompC* Δ*ompF* double mutant complemented in trans with either *ompC* or *ompF*. Interestingly, removing both OmpC and OmpF completely shifted the MlaA^K184C^ population to a MTSES-accessible state (**Fig. 5E**), and expressing OmpC or OmpF in trans (at comparable levels, **Fig. S7**) restored the population of MlaA^K184C^ to liken that in wild-type cells. These results indicate that free MlaA may not adopt a native conformational state in the OM, and that interactions with either porin is required yet sufficient for scaffolding MlaA. We further noted that the MlaA^K184C^ population in just the Δ*ompC* mutant was already fully MTSES-accessible, and this observation was independent of the presence of MlaC (**Fig. 5F**). Overall, MlaA in strains lacking OmpC alone may also mostly not be bound to porins (owing to low levels of OmpF), and therefore largely adopts a non-functional conformational state, mirroring what we have observed in the MlaA variants. We conclude that interactions with trimeric porins allow MlaA to adopt functional states important for lipid transport.

### Cryo-EM structure of the OmpC-MlaA^Q205C^-MlaC^V171C^ complex in nanodiscs provides a molecular picture for MlaC recruitment

To directly visualize possible conformational changes in MlaA upon MlaC binding, we sought to solve the complex structure of OmpC-MlaA bound to MlaC in a native lipid environment. We exploited the strong in vivo disulfide crosslinking to trap transient interactions between MlaA^Q205C^ and MlaC^V171C^ and effectively isolated such OmpC-MlaA-MlaC complexes from overexpressing cells, followed by purification on size-exclusion chromatography (SEC) (**Fig. S8A**). Multi-angle light scattering (MALS) analysis revealed on average two MlaA^Q205C^-MlaC^V171C^ bound to a trimer of OmpC in our purified detergent preparations (**Fig. S8B**). We reconstituted these OmpC-MlaA-MlaC complexes into lipid nanodiscs assembled using the MSP2N2 membrane scaffold protein (Denisov & Sligar, 2016) and *E. coli* polar lipids (**Fig. S9**) for single particle cryo-electron microscopy (cryo-EM) analysis.

Initial 3D reconstruction without application of symmetry operators revealed three distinct classes of OmpC-MlaA-MlaC complexes, varying in the number of MlaA-MlaC pairs bound per OmpC trimer (i.e. OmpC_3_-(MlaA-MlaC)_1-3_) (**Fig. S10**) (Punjani, Rubinstein, Fleet, & Brubaker, 2017). However, weak map densities suggested low occupancies for the additional MlaA-MlaC pairs (**Figs. S10 and S11**). Consequently, combining particles from these classes yielded an improved density map of OmpC_3_-(MlaA-MlaC) with an overall resolution of 2.9 Å (**Fig. S10A and S10C**).

The membrane-embedded components of OmpC3-MlaA were well-resolved with side chains clearly visualized, while the density of bound MlaC was limited to 5.0-7.0 Å (**Fig. 6A**) owing to continuous flexibility (Punjani & Fleet, 2021). The overall conformations of the OmpC trimer and MlaA in our complex (**Fig. 6B**) largely resembled those in reported crystal structures of *E. coli* OmpC alone (PDB 2J1N) and MlaA in complex with porins (PDB 5NUO-5NUR), respectively (**Fig. S12**). While MlaA binds the OmpC trimer at the porin subunit interface in a similar arrangement to that of porin-MlaA complexes (Abellon-Ruiz et al., 2017; Yeow et al., 2018), we did observe a slight downward tilt (~7 degrees) of MlaA pivoted at this contact site (PDB 5NUP) (**Fig. 6C**). This causes MlaA to lean towards the ‘periplasmic side’ of the nanodisc, which could either be due to the presence of a lipid bilayer and/or bound MlaC. In our nanodisc structure, we can clearly visualize the MlaA donut positioned exactly at the membrane-water boundary of the inner leaflet of the bilayer, with the C-terminal part of the protein protruding into periplasmic space and connected to MlaC at two contact points (**Fig. 6C**). Unbiased rigid body fitting of the MlaC model alone in the density positioned V171C adjacent to Q205C (on MlaA), exactly consistent with the engineered disulfide bond forming one of the contact points (**Fig. 6C**). In addition, we were able to trace and fit a poly-alanine model of the C-terminal tail helix of MlaA into residual density not occupied by MlaC, accounting for the second contact point. In the final real-space refined model, the manner by which the C-terminal helix of MlaA interacts with MlaC agrees with the AlphaFold2 prediction and strongly validates our biochemical observations (**Fig. 4**), solidifying the idea that this interaction is critical for MlaC recruitment. Unfortunately, it appears that we have captured MlaA-MlaC in a post-recruitment, pre-docking state, presumably constrained by the engineered disulfide linkage; that MlaC is not fully docked to the base of MlaA precluded observation of possible conformational changes expected during lipid transfer.

**Figure 6.**
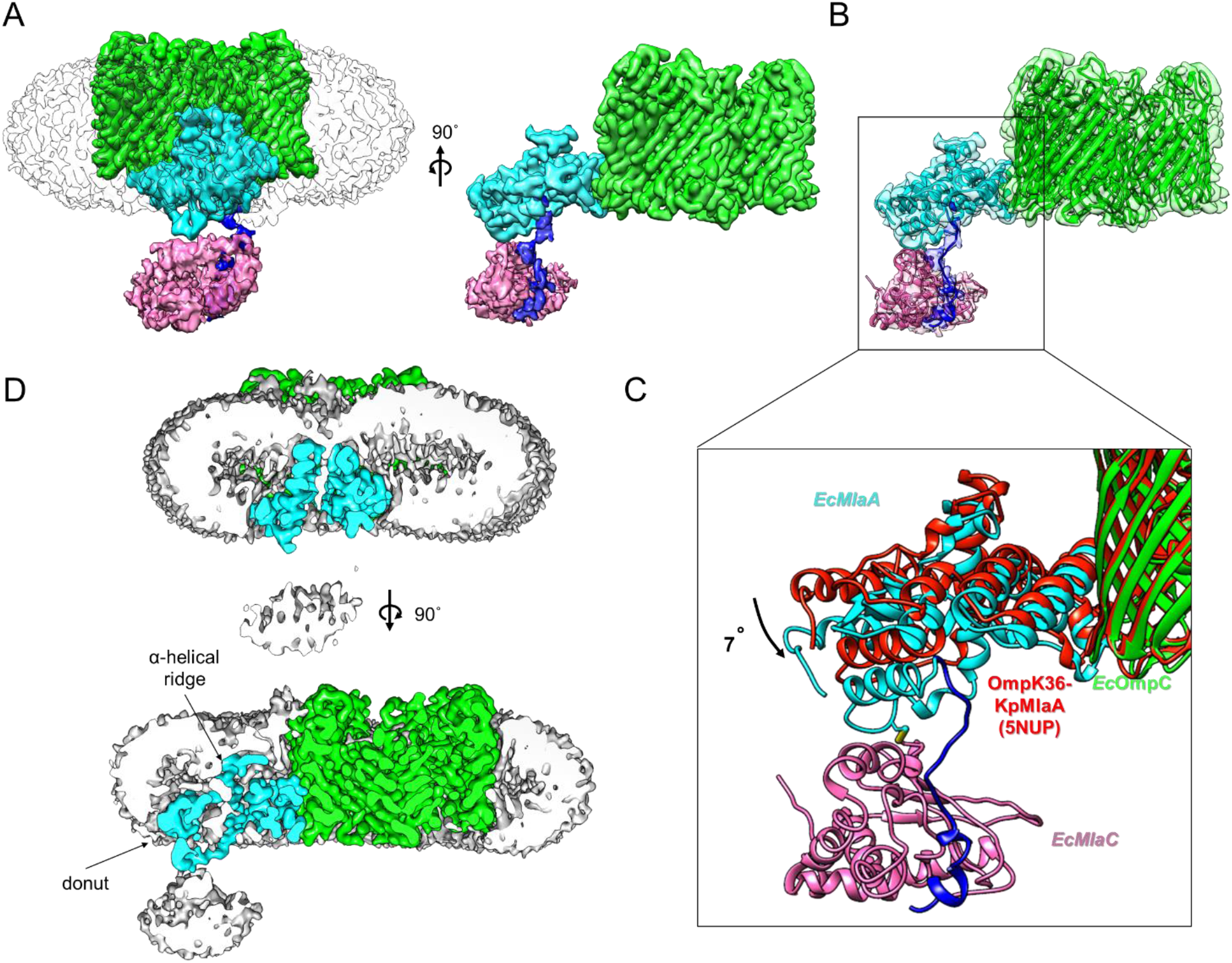
Cryo-EM structure of OmpC_3_-MlaA^Q205C^-MlaC^V171C^ in nanodiscs. **(A**) Front and side orientations of the density map of OmpC_3_-(MlaA-MlaC) (**EMD-35253**) (unsharpened; contour level of 0.06, *transparency 80%*) with the protein surface densities colored *green* (OmpC; contour level of 0.1), *cyan* (MlaA; contour level of 0.1), *blue* (C-terminal tail helix of MlaA; contour level of 0.06), and *pink* (MlaC; contour level of 0.06), respectively. **(B)**Cartoon illustrations of the OmpC_3_-MlaA-MlaC structure (**PDB 8I8X**) well-fitted and refined within protein surface densities in **(A)**(*transparency 80%*). Two contact points between MlaA and MlaC are clearly revealed (see **(C)**inset), one modelled with the engineered MlaA^Q205C^-MlaC^V171C^ disulfide bond, and the other with the C-terminal linker and tail helix of MlaA. **(C)**Superimposition of OmpC3-MlaA-MlaC (**PDB 8I8X**) and OmpK36-*Kp*MlaA (PDB 5NUP, *red*) structures revealed a 7° tilt of MlaA towards the periplasm. **(D)**Cut-away views of the density maps from **(A)**where the unsharpened nanodisc surface density is now colored *gray* (*transparency* 0%) revealed localized membrane thinning in the outer leaflet of the bilayer all around the position of the α-helical ridge of MlaA. Illustrations were generated using UCSF Chimera (Pettersen et al., 2004).

Regardless, our detailed molecular model of the complex, specifically MlaA, in a lipid bilayer enabled us to draw important mechanistic insights for the transfer reaction. While the MlaA donut sits in the inner leaflet of the bilayer, the channel ridge only protrudes midway into the outer leaflet. A really striking consequence of this placement is the mismatch between its membrane-spanning region and the thickness of the bilayer, leading to local membrane ‘thinning’, where the outer leaflet membrane-water boundary prominently bends inwards all around the position of the ridge (**Fig. 6D**). This deformation essentially creates a ‘funnel’ that likely selects for and guides outer leaflet PLs (headgroup first) into the hydrophilic channel of MlaA. Our structure of OmpC_3_-MlaA in a bilayer therefore reveals a possible mechanism for the initial steps leading to the removal of PLs from the outer leaflet of the OM.

## Discussion

How the OmpC-MlaA complex transfers PLs from the outer leaflet of the OM to MlaC, as part of the role of the OmpC-Mla system in maintaining lipid asymmetry, is not known. In this study, we have mapped the interacting surfaces between MlaA and MlaC, demonstrated conformational changes in MlaA during PL transfer to MlaC, and resolved the molecular architecture of an engineered OmpC_3_-MlaA-MlaC complex in a membrane bilayer. Using molecular cross-linking and computational modelling, we have shown that MlaC docks at the periplasmic face of MlaA, possibly in a manner where the lipid binding cavity of MlaC is aligned with the hydrophilic channel of MlaA to create a continuous path for PL movement. Furthermore, we have demonstrated that electrostatic interactions between the C-terminal tail helix of MlaA and a charged patch on MlaC are required for initial recruitment. We have also uncovered that wild-type MlaA adopts distinct conformational states that is likely modulated by interactions with (apo)-MlaC. Finally, we have elucidated the cryo-EM structure of nanodisc-embedded OmpC_3_-MlaA-MlaC trapped in a pre-docking state, revealing features that substantiated the mechanism for MlaC recruitment. While structural information on MlaA conformational changes was absent, we discerned a unique localized thinning effect on the bilayer by MlaA, pointing towards a possible way by which PLs may be funnelled into the MlaA channel. Our work provides critical mechanistic insights for the PL transfer reaction from OmpC-MlaA to MlaC for onward shuttling to the IM.

The requirement of electrostatic interactions between MlaA and MlaC allows us to infer a mechanism for how MlaA ensures effective recruitment and binding of apo-over holo-MlaC, thereby facilitating overall retrograde transfer of PLs. We have established that the positively-charged patch formed by residues K84/R90 on MlaC and the negatively-charged C-terminal tail helix residues D243/D244 on MlaA directly interact (**Fig. 4**), consistent with our structural observations (**Fig. 6**), and are important for MlaC recruitment. We note that residues K84/R90 on MlaC are exposed and accessible on apo-MlaC (Hughes et al., 2019). Remarkably, this positively-charged patch is somewhat occluded by movement of a surface loop on MlaC upon PL binding (Ekiert et al., 2017) (**Fig. 7**), limiting access by the MlaA tail helix. Such differential accessibility on MlaC likely allows preferential recruitment of apo-MlaC over holo-MlaC at the OM (**Fig. 7**). Post-recruitment, additional interactions between MlaA and MlaC occur at the base of the MlaA hydrophilic channel, stabilizing the complex for productive PL transfer. Once MlaC picks up a PL molecule, however, the charge-charge interaction with the MlaA tail helix is lost due to steric hindrance (**Fig. 7**), leading to possible destabilization of the MlaA-MlaC complex, and release of holo-MlaC from the OM. In a way, the MlaA tail helix, possibly extended almost halfway into the periplasmic space via a flexible linker (25 residues, ~88 Å), serves both as an electrostatic bait for MlaC during recruitment, and also a latch that locks down the transient MlaA-MlaC complex during PL transfer. This could be a compelling strategy to favor apo-MlaC recruitment/binding, and may ensure that PL transfer takes place in a retrograde fashion.

**Figure 7.**
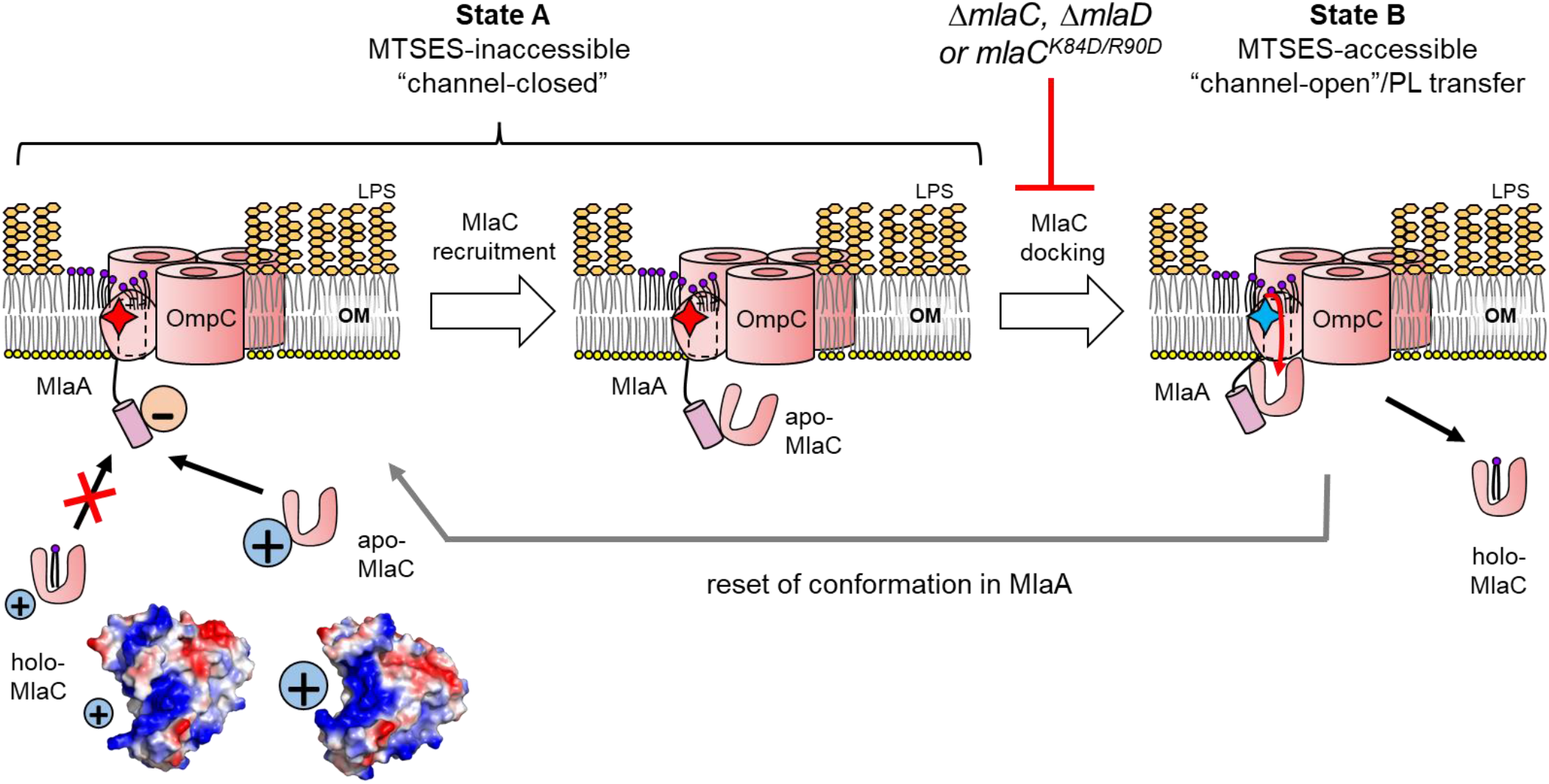
A proposed model for MlaC recruitment/binding and accompanying MlaA conformational changes that facilitate retrograde PL transfer by the OmpC-MlaA-(MlaC) complex. At the OM, apo-MlaC is preferentially recruited to MlaA via electrostatic interactions between MlaC surface charge patch and MlaA C-terminal tail helix. Subsequent docking of MlaC to the base of the MlaA channel induces a conformational change in MlaA, from an MTSES-<u>inaccessible</u> state (**State A**, “channel-closed”, residue K184 represented by *red star*) to an MTSES-<u>accessible</u> state (**State B**, “channel-open”, residue K184 represented by *blue star*) Along with the localized membrane thinning caused by MlaA, this conformational change may then facilitate the transfer of PLs from the outer leaflet of the OM into the lipid binding cavity of MlaC. Holo-MlaC then leaves the OmpC-MlaA complex to shuttle the PL ligand to the IM, thus resetting MlaA conformation. Vacuum electrostatic surface representations of MlaC bound with lipids (PDB 5UWA) (Ekiert et al., 2017), and in the unliganded form (PDB 6GKI) (Hughes et al., 2019).

In the context of MlaC binding and release, we can now also introduce a model for MlaA cycling between distinct conformation states during PL transfer (**Fig. 7**). We have shown that MlaA exists in at least two states in wildtype strains, where K184 is somehow accessible to the aqueous environment (**State B**), or not (**State A**). Interestingly, K184 is positioned midway up the MlaA channel in the membrane, and close to the gating loop; we therefore speculate that the MlaA channel may be open in **State B**, yet closed in **State A**. In the absence of MlaC, or with disrupted MlaA-MlaC recruitment, all of MlaA appears to become confined to **State A**(channel closed). Accordingly, apo-MlaC recruitment and binding is likely required to induce MlaA to shift to the channel-open **State B**, allowing productive PL transfer. Conceivably, holo-MlaC then no longer binds MlaA in quite the same way as apo-MlaC, possibly explaining why MlaA also adopts a channel-closed **State A** in the absence of MlaD, where all of MlaC is presumably in the holo-form. Such coordination between apo-MlaC binding and conformational changes in wild-type MlaA may favor retrograde PL transfer at the OM.

Conformational analysis using K184 accessibility also facilitated our understanding of non-functional variants/states of MlaA. We have found that free MlaA (not in complex with porins), MlaA^3G3P^ and MlaA* variants all adopt conformation state(s) where K184 is fully solvent accessible and is no longer responsive to the presence of MlaC (binding) (**Fig. 5**). Since K184 accessibility only produces binary output, however, free MlaA and these variants may well have distinct conformations that still allow K184 to be accessible to the aqueous environment. MlaA^3G3P^ is believed to be a loop-locked non-functional mutant, while MlaA* is thought to have lost control of its gating loop (Yeow et al., 2018). Our study suggests that even though MlaA* has gain-of-function possibly in allowing PLs to flip across the OM to the cell surface (Grimm et al., 2020; Sutterlin et al., 2016), this variant is in fact non-functional in the native role of MlaA. Consistent with this idea, it has recently been demonstrated that MlaA* is unable to support retrograde PL transfer in vitro (Tang et al., 2021). We have also now demonstrated that MlaA requires trimeric porins like OmpC and OmpF for scaffolding and possibly achieving functional conformations, thus overall accounting for porin function in the Mla system (Chong et al., 2015). Curiously, we did not observe changes in K184 accessibility in a strain expressing OmpC^R92A^ (**Fig. 5E**), a mutation at the subunit interface of OmpC trimers (where MlaA binds) that impacts OM lipid asymmetry (Yeow et al., 2018). We speculate that the extracellular loops, proximal to R92 and overlying MlaA, at the porin subunit interface could play additional roles, perhaps in helping to enrich mislocalized PLs for MlaA to gain easy access for removal. Of note, LPS tend to be bound at these porin subunit interfacial sites, as previously reported in OmpF X-ray crystal structures (Arunmanee et al., 2016). We have also detected densities at the same sites in our OmpC_3_-MlaA structure that can be confidently modelled as truncated LPS (Kdo_2_-Lipid A) molecules (**Fig. S13**). Interestingly, these LPS molecules appear to block possible site-specific interactions between OmpC and MlaA in the outer leaflet of the OM, as previously revealed by photocrosslinking studies (Yeow et al., 2018). We therefore propose that in the event of perturbed OM lipid asymmetry, bound LPS might be exchanged for PLs, allowing the latter to gain easy access to the MlaA channel through the membrane thinning/funnelling effect. Removing outer leaflet PLs from porin subunit interfaces eventually restores such porin-LPS, which has been implicated recently in the formation of ordered OMP lattices on the cell surface (Webby et al., 2022). Beyond proper scaffolding of MlaA, we believe that OmpC plays yet-to-be-ascertained functions that facilitate PL transport by MlaA.

It is truly fascinating that the manner by which MlaA sits in the bilayer causes localized membrane thinning. Such membrane perturbations/deformations have in fact been documented across protein machines that mediate the insertion and/or removal of substrates from a lipid bilayer (Botos, Noinaj, & Buchanan, 2017; Liu & Gumbart, 2020; Stubenrauch & Lithgow, 2019) (**Fig. 7**). We hypothesize that forcing outer leaflet lipids to bend towards its ridge structure offers a mechanism to favor PLs with flexible acyl tails, yet exclude highly rigid LPS molecules from, entering the MlaA channel. Furthermore, this effect facilitates the destabilization of individual PL molecules in the outer leaflet environment by reducing packing/non-covalent interactions with neighboring lipids, thereby enabling spontaneous and affinity-driven PL transfer to MlaC. MlaA is thus a uniquely evolved OM lipoprotein that catalyzes lipid gymnastics and movement. Ultimately, the exact mechanism by which the OmpC-MlaA complex extracts mislocalized PLs from the outer leaflet of the OM and transfers them to MlaC will require additional resolution into the various conformational states we have revealed herein. Structural information of OmpC-MlaA complexes with MlaC fully docked, as well as those of mutant MlaA variants, would provide further mechanistic insights into the process. Given that OM lipid asymmetry is critical for overall barrier function, novel insights into the OmpC-MlaA-(MlaC) complex will also guide us in developing useful strategies to inhibit the OmpC-Mla pathway in the future.

## Acknowledgements

J.Y. was supported by the National University of Singapore Graduate School of Integrative Sciences and Engineering Scholarship (ISEP). This work was supported by the Singapore Ministry of Health National Medical Research Council under its Open Fund Individual Research Grant (MOH-000145) (to S.S.C.). The authors would also like to acknowledge Dr Jian SHI from the Centre for Bio-Imaging Sciences (CBIS) at the National University of Singapore (NUS) for training and microscope facility management support. We also thank Dr Jianwei LI from Department of Biological Sciences (DBS), NUS for guidance with modelling and refinement techniques.

## Data Availability

Four 3D cryo-EM maps of OmpC_3_-(MlaA-MlaC) have been deposited in the Electron Microscopy Data Bank under accession numbers **EMD-35250** (OmpC_3_-(MlaA-MlaC)_1-3_), **EMD-35251** (OmpC_3_-(MlaA-MlaC)_3_), **EMD-35252** (OmpC_3_-(MlaA-MlaC)_2_), **EMD-35253** (OmpC_3_-(MlaA-MlaC)). Two atomic coordinate files have also been deposited in the Protein Data Bank under the accession numbers **8I8R** (OmpC_3_-MlaA) and **8I8X** (OmpC_3_-MlaA-MlaC).

