## Supplementary information for "Molecular mechanism of phospholipid transport at the bacterial outer membrane interface"

**This PDF file includes:**

Figures S1 to S13

Tables S1 to S4

SI References

**Supplemental Figures**

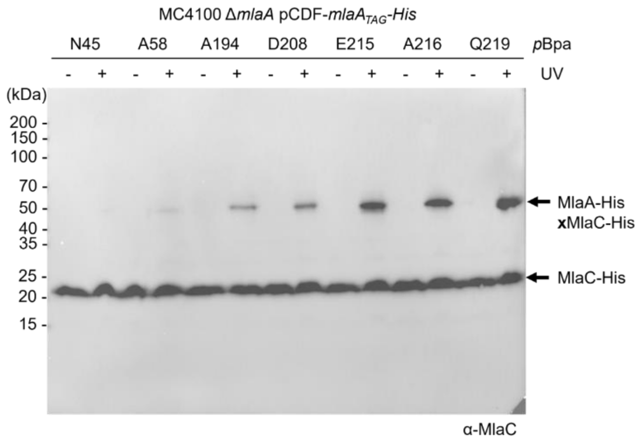


**Fig. S1.** Five more positions on the C-terminal α-helices of MlaA at the base of the hydrophilic channel interact with MlaC. Representative immunoblots showing UV-dependent formation of weak crosslinks between MlaA and MlaC in Δ*mlaA* cells expressing MlaA substituted with *p*Bpa at indicated positions from the pCDF plasmid.

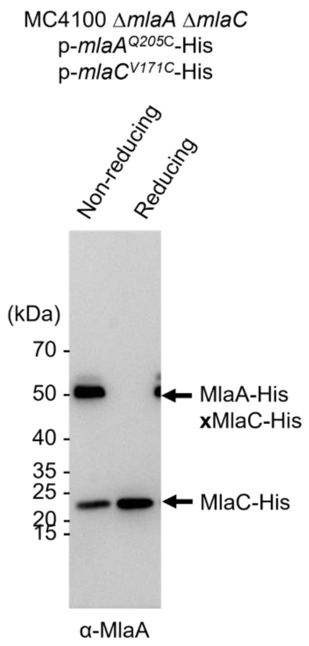


**Fig. S2.** Residue V171 at the opening of the lipid binding cavity of MlaC forms disulfide bonds extensively with residue Q205 at the periplasmic base of MlaA. Representative immunoblots showing formation of disulfide crosslinks between MlaA$^{Q205C}$ and MlaC$^{V171C}$ in Δ*mlaA* Δ*mlaC* cells expressing cysteine-substituted MlaA-His and MlaC-His from the pCDF and pET22/42 plasmids, respectively. The crosslinked band can be detected by α-MlaA (here) and α-MlaC (Fig. 2A). Samples were subjected to non-reducing or reducing SDS-PAGE prior to immunoblotting.

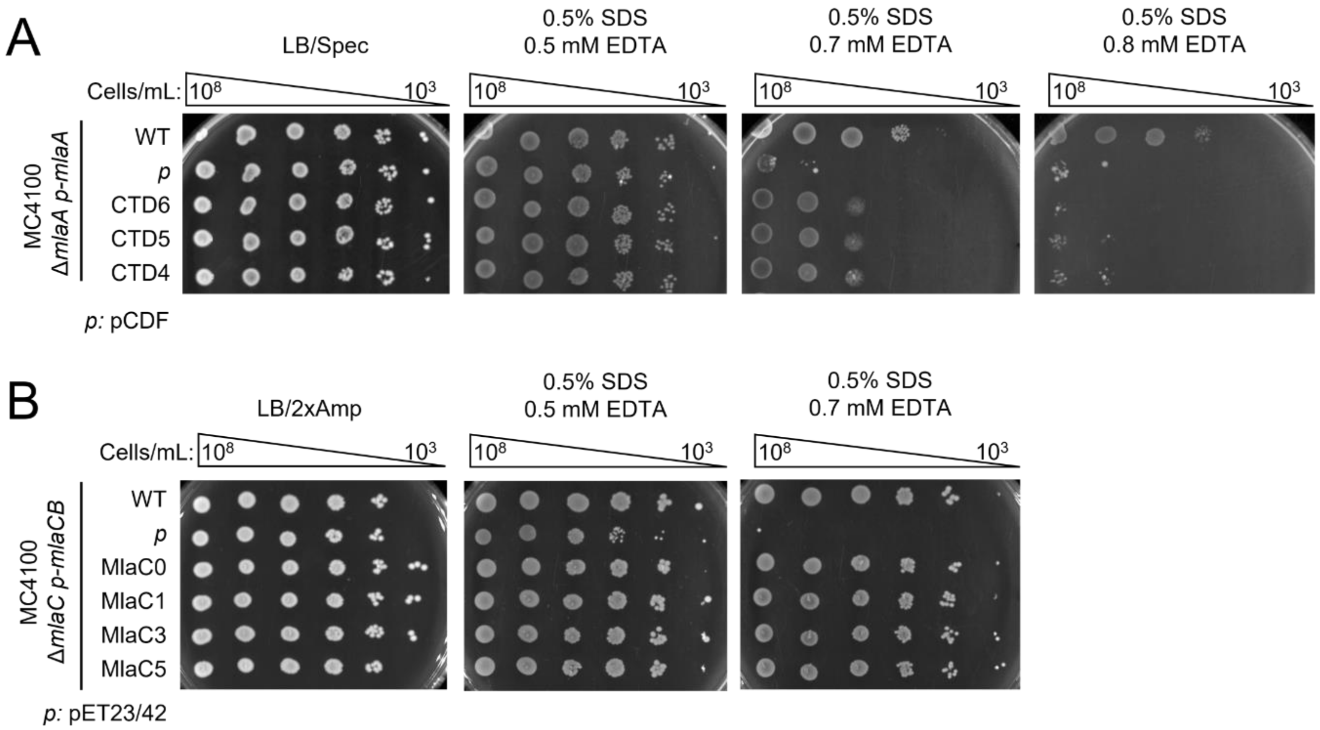


**Fig. S3.** Deletion of C-terminal tail helix of MlaA mildly perturbs Mla function, but charge-reversals at surface charge patches on MlaC do not. Analyses of SDS/EDTA sensitivity of (A) Δ*mlaA* strains producing indicated MlaA C-terminal tail helix deletion mutants from the pCDF vector, and (B) Δ*mlaC* strains producing indicated MlaC surface charge-reversed mutant variants from the pET23/42 vector.

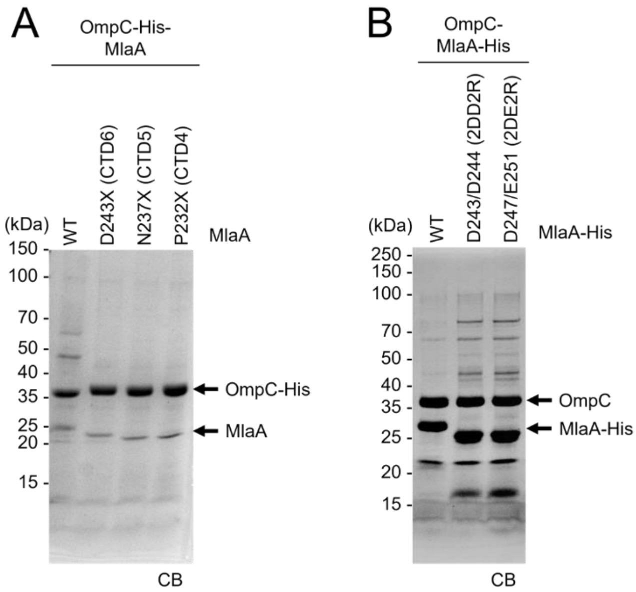


**Fig. S4.** MlaA C-terminal tail helix mutations do not affect OmpC-MlaA complex formation. Purified (A) (OmpC-His)-MlaA with indicated MlaA C-terminal tail helix deletion variants, and (B) OmpC-(MlaA-His) with indicated MlaA C-terminal tail helix charge-reversed variants subjected to SDS-PAGE, followed by Coomassie blue (CB) staining.

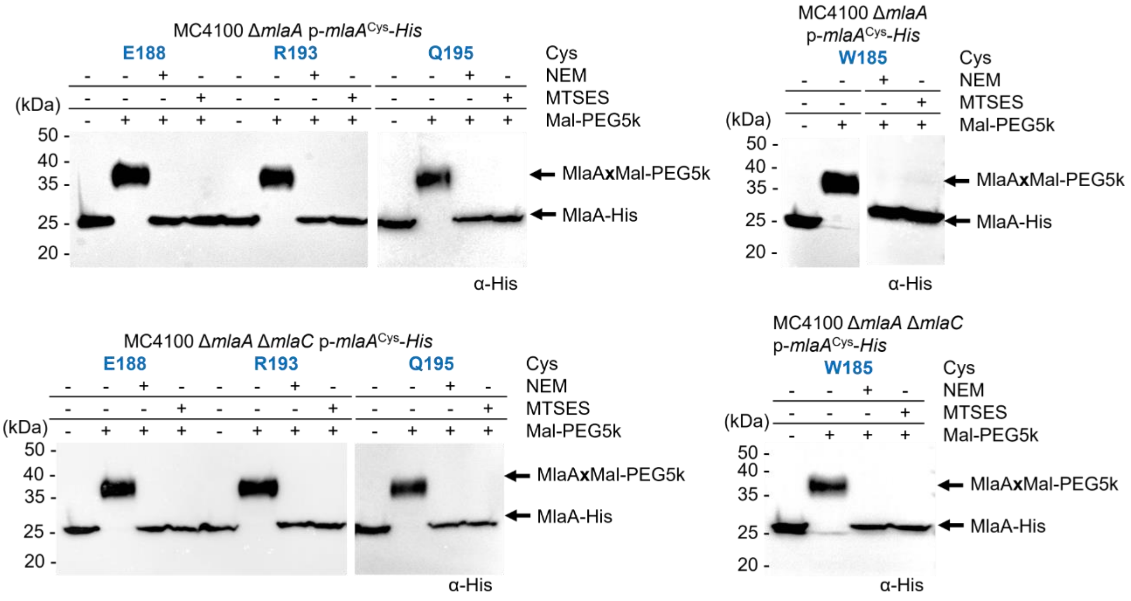


**Fig. S5.** Other selected MlaA channel residues do not display solvent accessibility changes in the absence of MlaC. Representative immunoblots showing Mal-PEG5k alkylation of MlaA variants containing channel residues substituted with cysteine (in Δ*mlaA* or Δ*mlaA* Δ*mlaC* strains) following labelling by membrane permeable N-ethylmaleimide (NEM) or impermeable (MTSES) reagents. Mal-PEG5k alkylated $MlaA^{Cys}$-His variants show an approximate ~5 kDa mass shift. Positions fully blocked by MTSES, which reflects the level of solvent accessibility, are highlighted in *blue*.

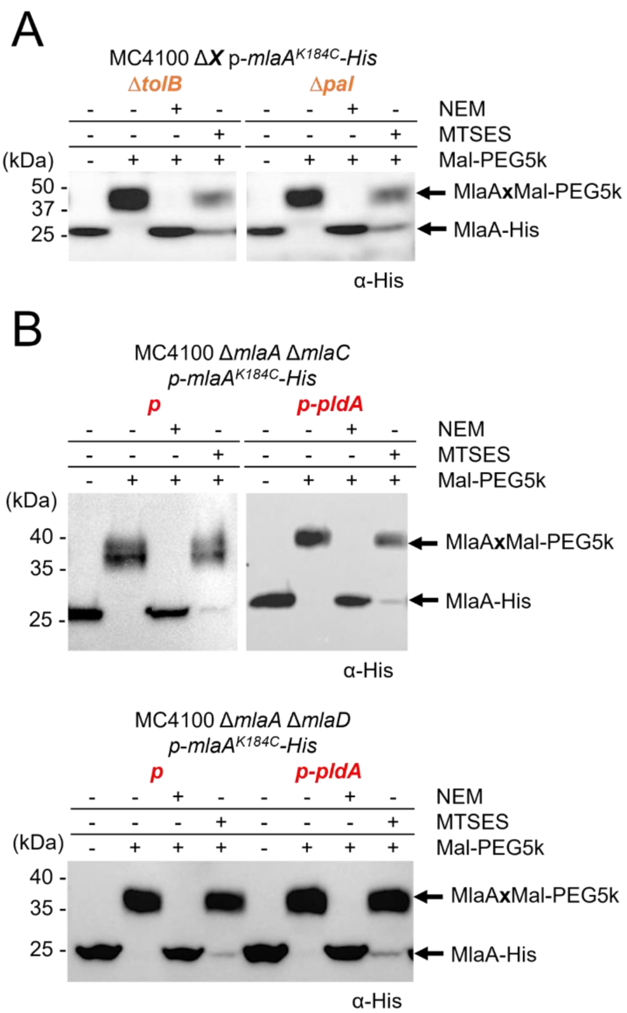


**Fig. S6.** MlaA channel solvent accessibility changes at residue K184 is not influenced by perturbation of OM lipid asymmetry. Representative immunoblots showing Mal-PEG5k alkylation of MlaA$^{K184C}$-His variant expressed from the pCDF plasmid, either in **(A)** background strains harboring disrupted lipid asymmetry (i.e. Δ*tolB* and Δ*pal*) or **(B)** in Δ*mlaA* Δ*mlaC* or Δ*mlaA* Δ*mlaD* strains also overproducing PldA from the pBR322 plasmid. Cells were labelled with membrane-permeable (NEM) or impermeable (MTSES) reagents, followed by alkylation with Mal-PEG5k, which introduces a ~5-kDa mass shift to MlaA$^{K184C}$-His. The levels of solvent accessibility of K184C in MlaA in the various strains, i.e. partially, or not blocked by MTSES, are highlighted in *orange* or *red*, respectively.

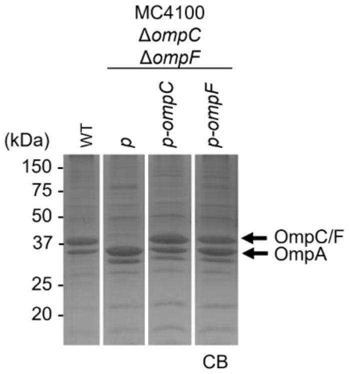


**Fig. S7.** OmpC and OmpF expressed in trans from the pDSW206 plasmid achieve native levels of porin expression (OmpC/F) in wild-type cells. Membrane fractions of wild-type or Δ*ompC* Δ*ompF* cells expressing OmpC or OmpF from pDSW206 plasmids were subjected to SDS-PAGE, followed by Coomassie blue (CB) staining.

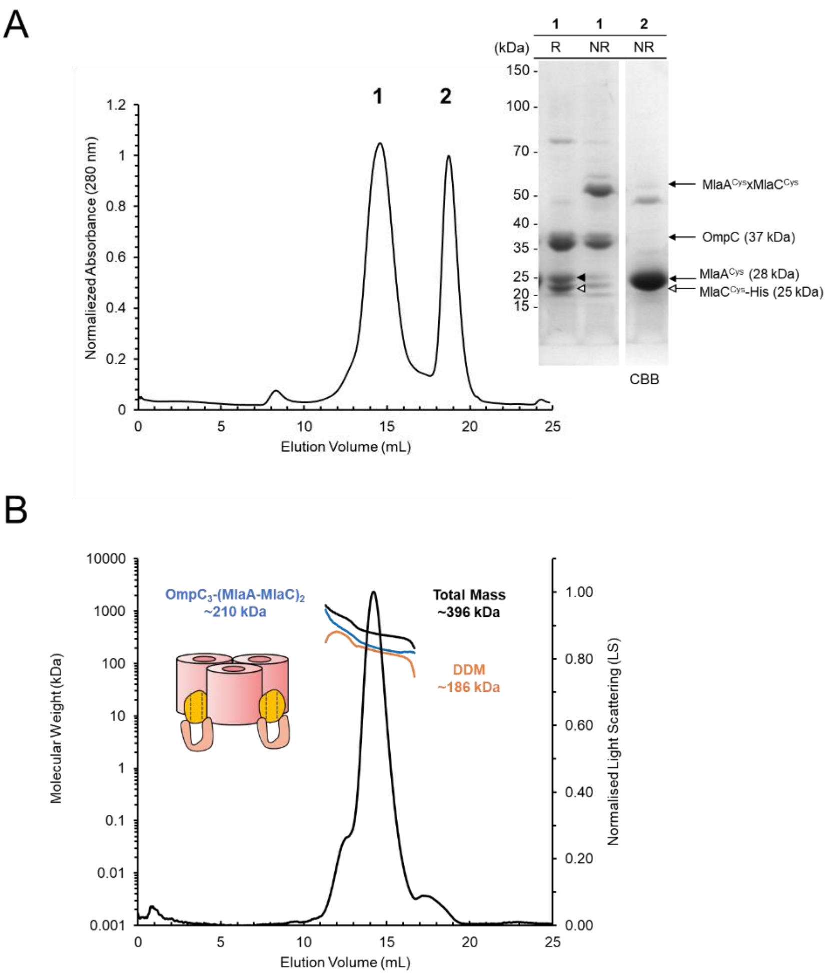


**Fig. S8.** The disulfide-trapped OmpC$_3$-MlaA$^{Q205C}$-MlaC$^{V171C}$ complexes can be purified to homogeneity. **(A)** Size exclusion chromatographic (SEC) analysis of OmpC$_3$-MlaA$^{Cys}$-MlaC$^{Cys}$-His in DDM. Peaks 1 and 2 correspond to OmpC$_3$-MlaA$^{Cys}$-MlaC$^{Cys}$-His and free MlaC$^{Cys}$-His, respectively. Comparative reducing (R) and non-reducing (NR) SDS-PAGE analyses of peak fractions (*right*) revealed the formation of disulfide crosslink between MlaA$^{Q205C}$ and MlaC$^{V171C}$ in our purified preparation. **(B)** SEC/multi-angle light scattering (MALS) analysis of OmpC$_3$-MlaA$^{Cys}$-MlaC$^{Cys}$-His in DDM. Molecular weight of OmpC$_3$-(MlaA$^{Cys}$-MlaC$^{Cys}$)$_2$ : ~213 kDa (predicted), ~210 (±2.2%) kDa (observed). Molecular weight of DDM fraction in the complex was ~186 (±5.7%) kDa. Numbers stated after ± show statistical consistency of analysis.

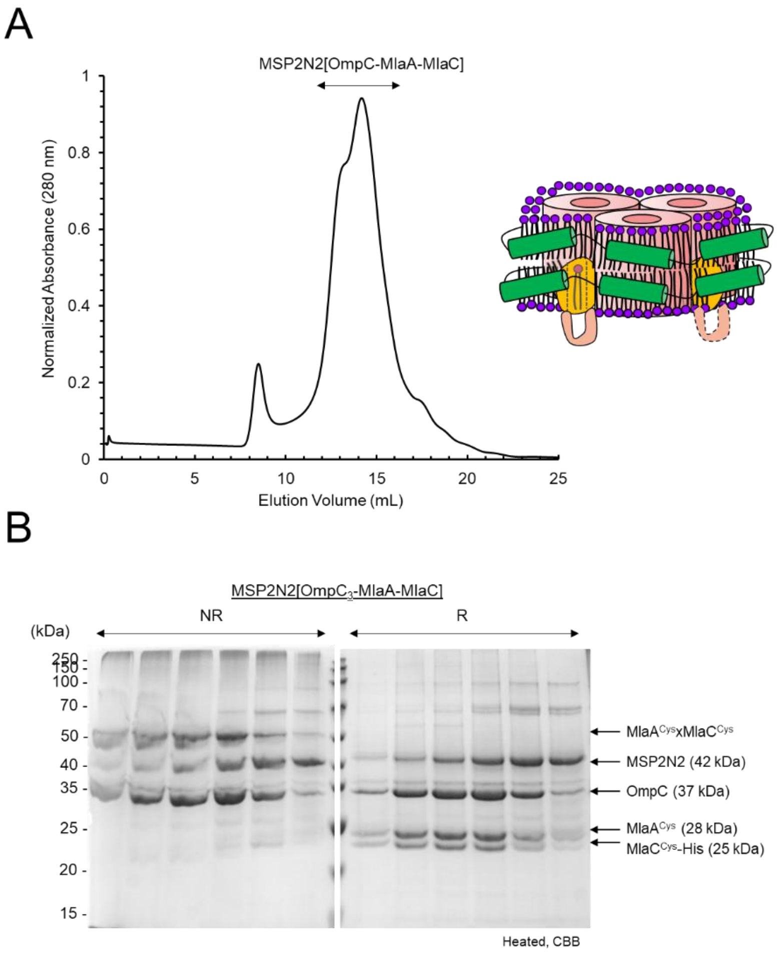


**Fig. S9.** Reconstitution of the $OmpC_3$-MlaA$^{Q205C}$-MlaC$^{V171C}$ complexes in nanodiscs. **(A)** SEC analysis of $OmpC_3$-MlaA$^{Cys}$-MlaC$^{Cys}$-His reconstituted in MSP2N2 nanodiscs. **(B)** Peak fractions of interest were subjected to reducing (R) and non-reducing (NR) SDS-PAGE analyses.

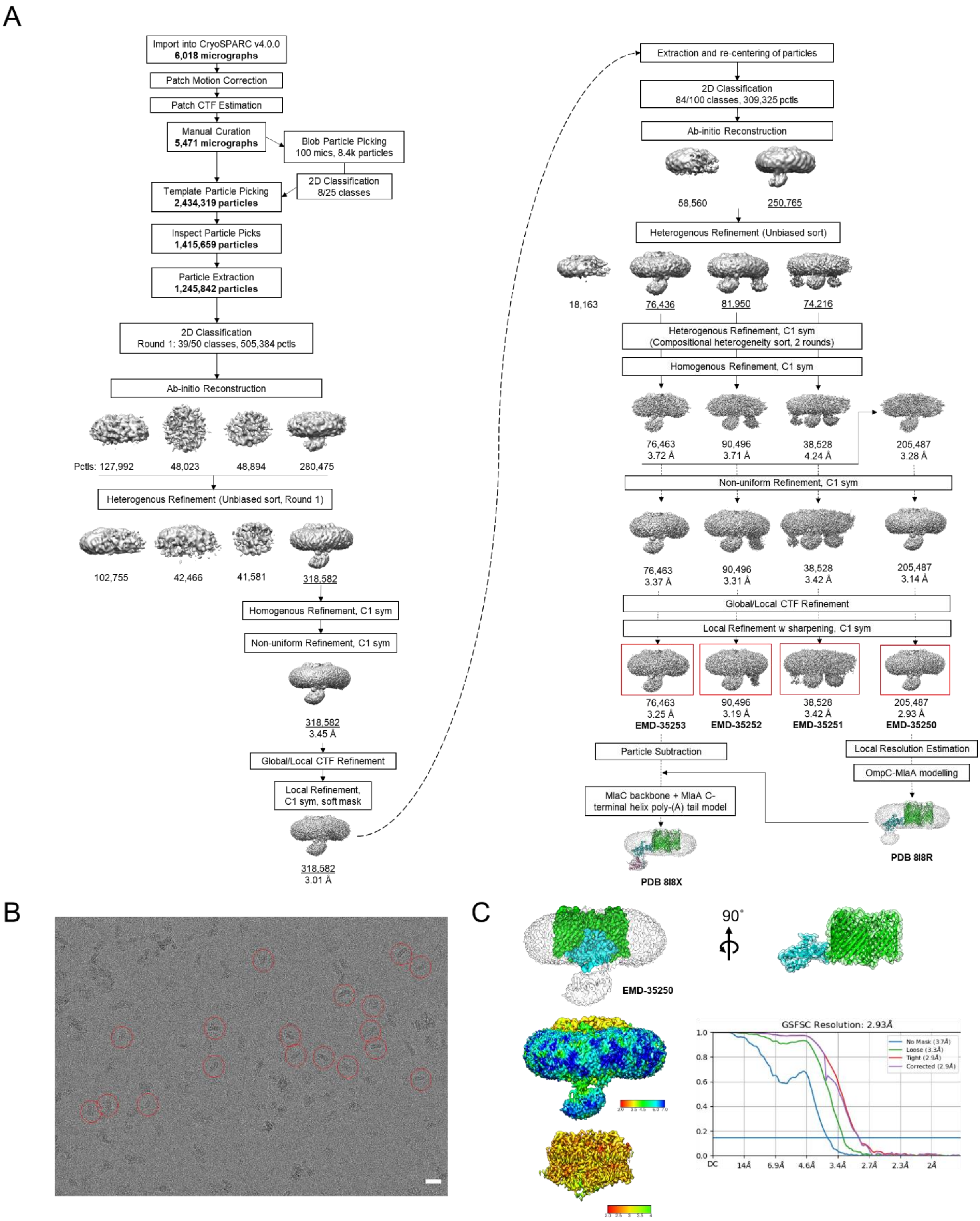
A
Import into CryoSPARC v4.0.0
6,018 micrographs
Patch Motion Correction
Patch CTF Estimation
Manual Curation
5,471 micrographs
Blob Particle Picking
100 mics, 8.4k particles
2D Classification
8/25 classes
Template Particle Picking
2,434,319 particles
Inspect Particle Picks
1,415,659 particles
Particle Extraction
1,245,842 particles
2D Classification
Round 1: 39/50 classes, 505,384 pctls
Ab-initio Reconstruction
Pctls: 127,992
48,023
48,894
280,475
Heterogenous Refinement (Unbiased sort, Round 1)
102,755
42,466
41,581
318,582
Homogenous Refinement, C1 sym
Non-uniform Refinement, C1 sym
318,582
3.45 Å
Global/Local CTF Refinement
Local Refinement,
C1 sym, soft mask
318,582
3.01 Å
Extraction and re-centering of particles
2D Classification
84/100 classes, 309,325 pctls
Ab-initio Reconstruction
58,560
250,765
Heterogenous Refinement (Unbiased sort)
18,163
76,436
81,950
74,216
Heterogenous Refinement, C1 sym
(Compositional heterogeneity sort, 2 rounds)
Homogenous Refinement, C1 sym
76,463
3.72 Å
90,496
3.71 Å
38,528
4.24 Å
205,487
3.28 Å
Non-uniform Refinement, C1 sym
76,463
3.37 Å
90,496
3.31 Å
38,528
3.42 Å
205,487
3.14 Å
Global/Local CTF Refinement
Local Refinement w sharpening, C1 sym
76,463
3.25 Å
EMD-35253
90,496
3.19 Å
EMD-35252
38,528
3.42 Å
EMD-35251
205,487
2.93 Å
EMD-35250
Particle Subtraction
Local Resolution Estimation
OmpC-MlaA modelling
MlaC backbone + MlaA C-terminal helix poly-(A) tail model
PDB 8I8X
PDB 8I8R
B
C
EMD-35250
90°
GSFSC Resolution: 2.93Å
No Mask (3.7Å)
Loose (3.3Å)
Tight (2.9Å)
Corrected (2.9Å)
1.0
0.8
0.6
0.4
0.2
0.0
DC
14Å
6.9Å
4.6Å
3.4Å
2.7Å
2.3Å
2Å

**Fig. S10.** Single-particle cryo-EM analysis of nanodisc-embedded $OmpC_3$- $MlaA^{Q205C}$-$MlaC^{V171C}$ complexes. **(A)** Data processing flowchart yielding the final maps (**EMD-35321/2/3**) of three major classes representing compositional heterogeneity in $OmpC_3$-$(MlaA\text{-}MlaC)_x$. A fourth map for $OmpC_3$-$(MlaA\text{-}MlaC)_{1\text{-}3}$ (**EMD-35250**) was generated, where all particles from the above classes were combined and refined. The structure for $OmpC_3$-MlaA (**PDB 8I8R**) was built and refined in **EMD-35250**, while MlaC and the MlaA C-terminal tail helix (**PDB 8I8X**) were modelled in **EMD-35253**. Additional refinement details can be found in **Table S4**. **(B)** Representative cryo-EM image with several particles marked by circles. **(C)** Front and side orientations of the density map of $OmpC_3$-$(MlaA\text{-}MlaC)_{1\text{-}3}$ (**EMD-35250**) (unsharpened; contour level of 0.06, *transparency 80%*) with the protein surface densities colored *green* (OmpC; contour level of 0.1) and *cyan* (MlaA; contour level of 0.1) according to **PDB 8I8R**. Unsharpened and sharpened local resolution maps were calculated by cryoSPARC (Punjani, Rubinstein, Fleet, & Brubaker, 2017), and illustrated with pseudo-color representation of per-voxel resolution. The gold standard-Fourier Shell Correlation (GS-FSC) plots of unmasked and masked (loose, tight, and corrected) maps are derived from cryoSPARC from local refinement. Illustrations were generated using the software UCSF Chimera (Pettersen et al., 2004).

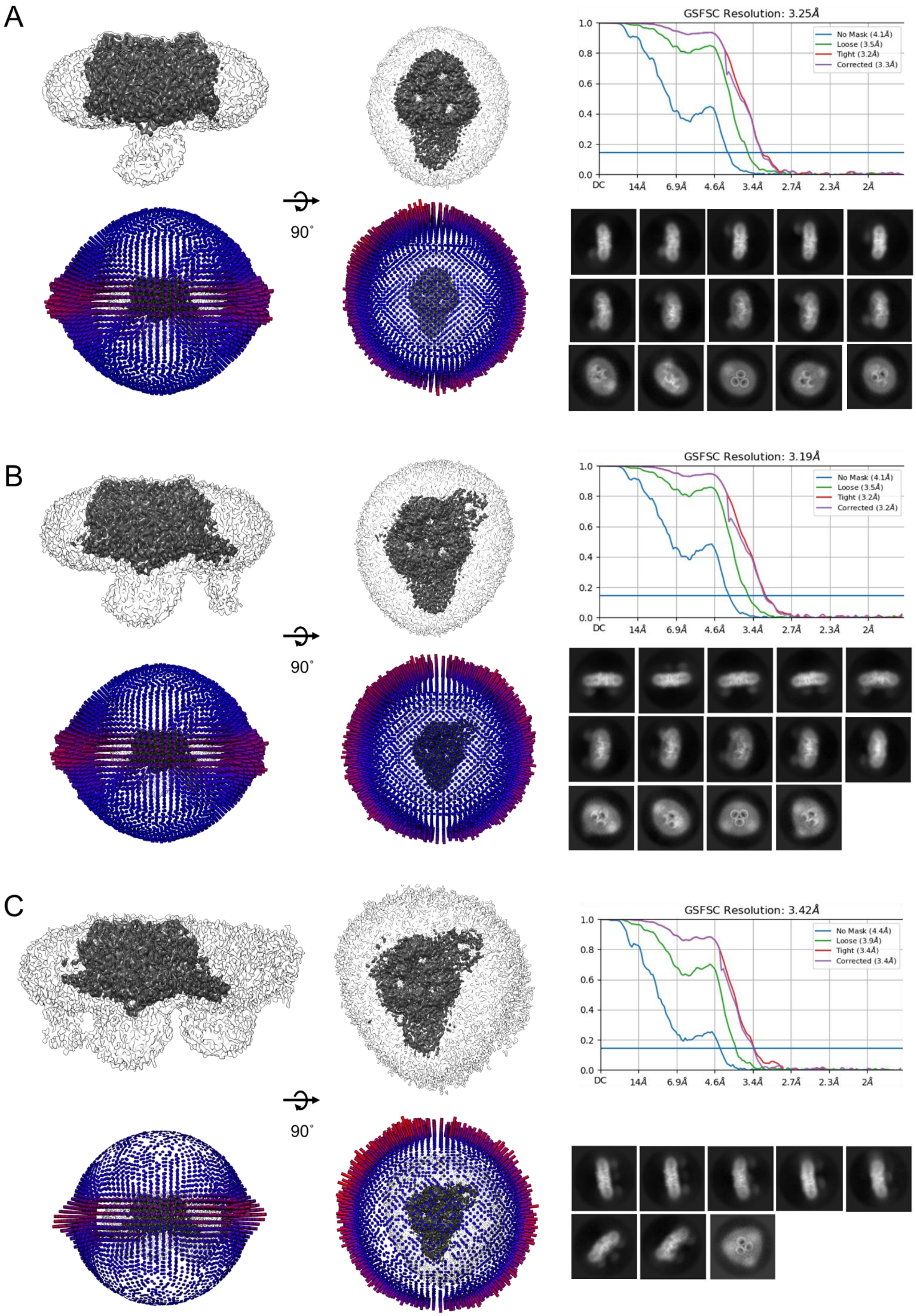
A
GSFSC Resolution: 3.25Å
No Mask (4.1Å)
Loose (3.5Å)
Tight (3.2Å)
Corrected (3.3Å)
1.0
0.8
0.6
0.4
0.2
0.0
DC
14Å
6.9Å
4.6Å
3.4Å
2.7Å
2.3Å
2Å
90˚
B
GSFSC Resolution: 3.19Å
No Mask (4.1Å)
Loose (3.5Å)
Tight (3.2Å)
Corrected (3.2Å)
90˚
C
GSFSC Resolution: 3.42Å
No Mask (4.4Å)
Loose (3.9Å)
Tight (3.4Å)
Corrected (3.4Å)
90˚

**Fig. S11.** Cryo-EM density maps of various compositional variants of $OmpC_3$-($MlaA^{Q205C}$-$MlaC^{V171C}$)$_x$ in nanodiscs. **(A-C)** Relevant maps (sharpened/unsharpened) and parameters (Euler angle distributions, GS-FSC plots, representative 2D classes) for **(A)** $OmpC_3$-(MlaA-MlaC) (**EMD-35253**), **(B)** $OmpC_3$-(MlaA-MlaC)$_2$ (**EMD-35252**) and **(C)** $OmpC_3$-(MlaA-MlaC)$_3$ (**EMD-35251**) are presented. Sharpened (contour levels of 0.11, coloured *gray*) and unsharpened (contour levels of 0.06, coloured *white, transparency 80%*) densities are shown in *front* and *top* orientations. For Euler angle distributions, the height and color of each rod is proportional to the amount of particles visualized from the same specific orientation. The gold standard-Fourier Shell Correlation (GS-FSC) plots of unmasked and masked (loose, tight, and corrected) maps are derived from cryoSPARC. Representative 2D classes are generated using a particle box size of 320 pixels (275 Å). Illustrations were generated using the software UCSF Chimera (Pettersen et al., 2004).

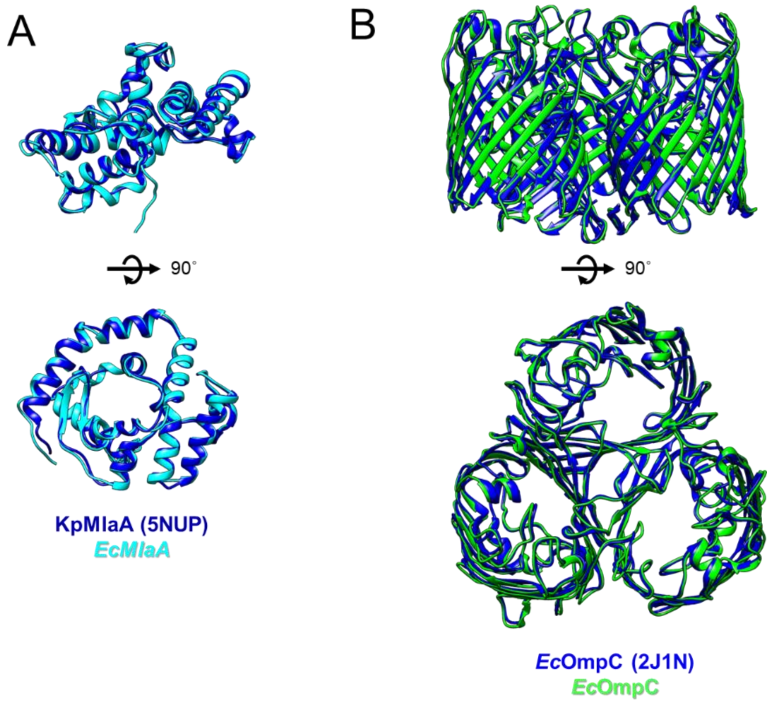


**Fig. S12.** Refined models of OmpC trimer and MlaA are highly similar to reported structures. Superimpositions of **(A)** OmpC trimer in our model (**PDB 8I8R**) with the reported OmpC trimer crystal structure (PDB 2J1N), and of **(B)** *Ec*MlaA (**PDB 8I8R**) and the reported *Kp*MlaA structure in OmpK36-*Kp*MlaA (PDB 5NUP) reveal insignificant root mean square deviations (r.m.s.d. <1 Å). Illustrations were generated using the software UCSF Chimera (Pettersen et al., 2004).

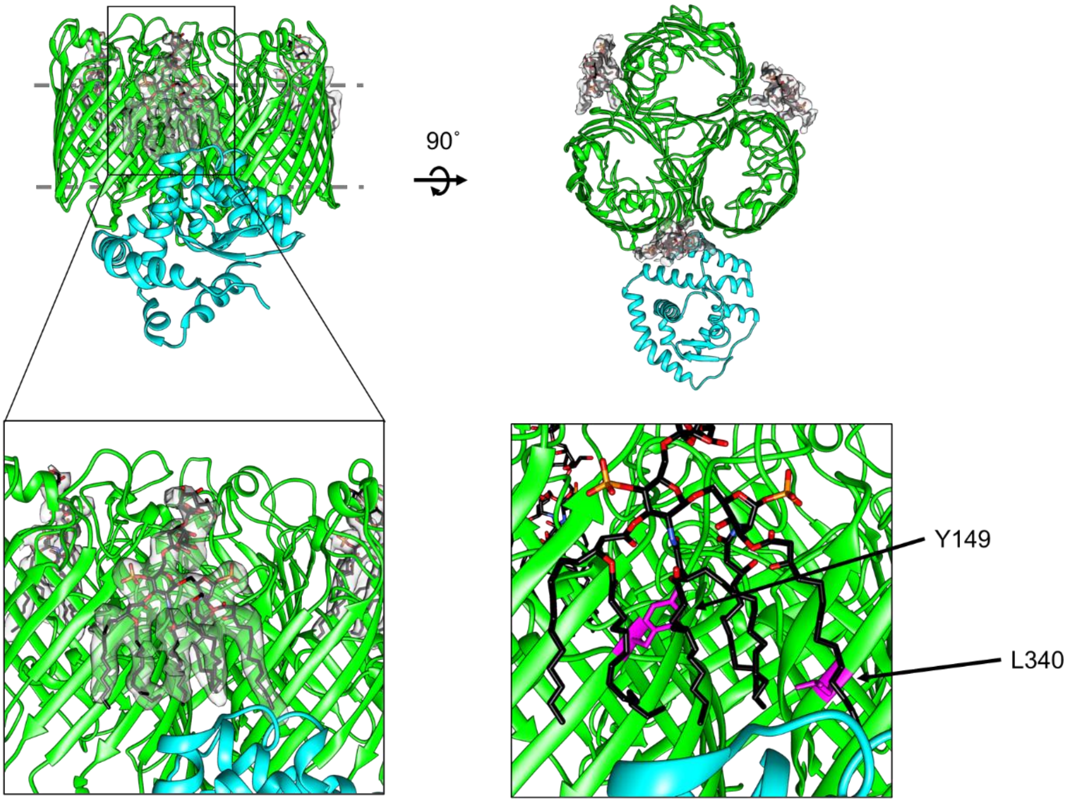


**Fig. S13.** LPS occupy interfaces of porin trimer subunits. Extra densities (*gray*; contour level 0.08) at porin trimer subunit interfaces in our OmpC$_3$-MlaA model (**PDB 8I8R**) can be confidently modelled as truncated LPS molecules (Kdo$_2$-Lipid A (KDL)). These LPS molecules occlude positions (Y149 and L340, *purple sticks*) on OmpC that enabled previously-reported photo-crosslinking to MlaA in the outer leaflet of the OM (Yeow et al., 2018). Cartoon illustrations are colored *green* (OmpC) and *cyan* (MlaA), respectively. Illustrations were generated using the software UCSF Chimera (Pettersen et al., 2004).

### Materials and Methods

**Bacterial strains and plasmids.** All strains, plasmids, and primers used are listed in Table S1, S2, and S3 respectively.

**Growth conditions.** Lysogeny Broth (LB) and agar were prepared as previously described (Bertani, 2004). Unless otherwise noted, ampicillin (Amp) (Sigma-Aldrich, MO, USA) was used at a concentration of 200 µg/mL, chloramphenicol (Cam) (Alfa Aesar, Heysham, UK) at 15 µg/mL, kanamycin (Kan) (Sigma-Aldrich) at 25 µg/mL, and spectinomycin (Spec) (Sigma-Aldrich) at 50 µg/mL. For crosslinking experiments, *para*-benzoyl-L-phenylalanine (*p*Bpa; Alfa Aesar) was dissolved in 1 M NaOH at 0.25 M, and used at 0.25 mM unless otherwise mentioned.

**In vivo photoactivable crosslinking.** We adopted previously described protocol (Chin, Martin, King, Wang, & Schultz, 2002) for all in vivo photoactivable crosslinking experiments. Briefly, amber stop codon (TAG) was introduced at selected positions in pCDF*mlaA-His* plasmids via site directed mutagenesis using primers listed in Table S3. For MlaA crosslinking, MC4100 with Δ*mlaA*::*kan* background harbouring p*Sup-BpaRS-6TRN* (Ryu & Schultz, 2006) and pCDF*mlaA*$_{TAG}$*-His* were used. An overnight 5 mL culture was grown from a single colony in LB broth supplemented with appropriate antibiotics at 37 °C. Overnight cultures were diluted 1:100 into 10 mL of the same media containing 0.25 mM *p*Bpa and grown until $OD_{600}$ reached ~1.0. Cells were normalized by optical density before pelleting and resuspended in 1 mL ice cold TBS (20 mM Tris pH 8.0, 150 mM NaCl). Samples were either used directly or irradiated with UV light at 365 nm for 20 min at 4 °C or room temperature. All samples were pelleted again and finally resuspended in 200 µL of 2 X Laemmli buffer, boiled for 10 min, and centrifuged at 21,000 x *g* in a microcentrifuge for one min at room temperature; 15 µL of each sample subjected to SDS-PAGE and immunoblot analyses.

**In vivo disulfide bond analysis.** Cells harbouring pCDF*mlaA*$_{Cys}$*-His* and pET22/42*mlaC*$_{Cys}$*-His* expressing MlaA$_{Cys}$-His and MlaC$_{Cys}$-His respectively with site specific cysteine substitutions was grown overnight supplemented with appropriate antibiotics in LB broth at 37 °C. Overnight cultures were diluted 1:100 into 5 mL of the same media and grown until $OD_{600}$ reached ~1.0. Cells were normalized by optical density. All samples were pelleted again and finally resuspended in 100 μL of mixed with 2 X Laemmlli buffer (reducing or non-reducing), boiled for 10 min and subjected to SDS-PAGE and immunoblotting analyses using α-MlaC antibody.

**Substituted cysteine accessibility method (SCAM).** For experiments assessing the function of MlaA mutants, cells harbouring pET23/42*mlaA$_{Cys}$-His* ($Amp^R$) expressing $MlaA_{Cys}$-His in various wildtype and *mla* background strains were grown overnight and supplemented with appropriate antibiotics in LB broth at 37 °C. To assess porin function, cells harbouring pDSW206-*ompC/F* ($Amp^R$) and pCDF*mlaA$_{Cys}$-His* ($Spec^R$) expressing $MlaA_{Cys}$-His and native levels of porins in Δ*ompC ompF* background were grown overnight and supplemented with appropriate antibiotics in LB broth at 37 °C. Overnight cultures were diluted 1:100 into 5 mL of the same media and grown until $OD_{600}$ reached ~1.0. Cells were normalized by optical density. 1-mL cells were grown to exponential phase ($OD_{600}$ ~0.6), washed twice with TBS (pH 8.0), and resuspended in 480 μL of TBS. For the blocking step, four tubes containing 120 μL of cell suspension were either untreated (positive and negative control tubes added with deionized $H_2O$) or treated with 5 mM thiol-reactive reagent *N*-ethylmaleimide (NEM, Thermo Scientific) or sodium (2-sulfonatoethyl) methanethiosulfonate (MTSES, Biotium). As MTSES is membrane impermeable, it is expected to react with the free cysteine in MlaA variants only when the residue near or at the membrane-water boundaries, or in a hydrophilic channel. In contrast, NEM is expected to label all MlaA cysteine variants as it is membrane permeable. Reaction with MTSES or NEM blocks the particular cysteine site from subsequently labelling by maleimide-polyethylene glycol (Mal-PEG; 5 kDa, Sigma-Aldrich). After agitation at room temperature for 1 h, cells were washed twice with TBS, pelleted at 16,000 x *g*, and resuspended in 100 μL of lysis buffer (10 M urea, 1% SDS, 2 mM EDTA in 1 M Tris pH 6.8). Both NEM- and MTSES-blocked samples and the positive control sample were exposed to 1.2 mM Mal-PEG-5k. After agitation for another hour with protection from light, all samples were added with 120 μL of 2 X Laemmli buffer, boiled for 10 min, and centrifuged at 21,000 x *g* in a microcentrifuge for one min at room temperature; 20 μL from each sample tubes were subjected to SDS-PAGE and immunoblot analyses.

**Over-expression and purification of OmpC-MlaA-His or OmpC-MlaA$^{Cys}$-MlaC$^{Cys}$-His complexes.** To examine interactions of MlaA mutants with OmpC, OmpC-MlaA-His protein complexes were over-expressed and purified from BL21(λDE3) Δ*ompF*::*kan* cells (Yeow et al., 2018) co-transformed with either pDSW206*ompC-His* and pCDF*mlaA*, or pDSW206*ompC* and pCDF*mlaA-His*. To obtain OmpC-MlaA$^{Cys}$-MlaC$^{Cys}$-His complexes for structural analysis, OmpC, MlaA$^{Q205C}$ and MlaC$^{V171C}$-His were over-expressed and purified from BL21(λDE3) Δ*ompF*::*kan* cells (Yeow et al., 2018) co-transformed with pDSW206*ompC* and pCDF*mlaC$^{Cys}$-His-mlaA$^{Cys}$*. An overnight 10-mL culture was grown from a single colony in LB broth supplemented with appropriate antibiotics at 37 °C. The cell culture was then used to inoculate a 1-L culture and grown at the same temperature until $OD_{600}$ reached ~ 0.6. For induction, 0.5 mM IPTG (Axil Scientific, Singapore) was added and the culture was grown for another 3 h at 37 °C. Cells were pelleted by centrifugation at 4,700 x *g* for 20 min and then resuspended in 10-mL TBS containing 1 mM PMSF (Calbiochem) and 30 mM imidazole (Sigma-Aldrich). Cells were lysed with three rounds of sonication on ice (38 % power, 1 second pulse on, 1 second pulse off for 3 min). Cell lysates were incubated overnight with 1 % n-dodecyl β-D-maltoside (DDM, Calbiochem) at 4 °C. Cell debris was removed by centrifugation at 24,000 x *g* for 30 min at 4 °C. Subsequently, supernatant was incubated with 1 mL Ni-NTA nickel resin (QIAGEN), pre-equilibrated with 20 mL of wash buffer (TBS containing 0.05 % DDM and 50 mM imidazole) in a column for 1 h at 4 °C with rocking. The mixture was allowed to drain by gravity before washing vigorously with 10 x 10 mL of wash buffer and eluted with 10 mL of elution buffer (TBS containing 0.05 % DDM and 500 mM imidazole). The eluate was concentrated in an Amicon Ultra 100 kDa cut-off ultra-filtration device (Merck Millipore) by centrifugation at 4,000 x *g* to ~500 μL. Proteins were further purified by size-exclusion chromatography (AKTA Pure, GE Healthcare, UK) at 4 °C on a prepacked Superose 6 increase 10/300 GL column, using TBS containing 0.05 % DDM as the eluent.

**SDS-PAGE, immunoblotting and staining.** All samples subjected to SDS-PAGE were mixed 1:1 with 2X Laemmli buffer. Except for temperature titration experiments, the samples were subsequently either kept at room temperature or subjected to boiling at 100 °C for 10 min. Equal volumes of the samples were loaded onto the gels. As indicated in the figure legends, SDS-PAGE was performed using either 12% Tris.HCl gels (Laemmli, 1970) at 200 V for 45 min. After SDS-PAGE, gels were visualized by either Coomassie Blue staining, or subjected to immunoblot analysis. Immunoblot analysis was performed by transferring protein bands from the gels onto polyvinylidene fluoride (PVDF) membranes (Immun-Blot 0.2 µm, Bio-Rad, CA, USA) using semi-dry electroblotting system (Trans-Blot Turbo Transfer System, Bio-Rad). Membranes were blocked for 1 h at room temperature by 1 X casein blocking buffer (Sigma-Aldrich), washed and incubated with either primary antibodies (monoclonal α-MlaA (Chong, Woo, & Chng, 2015) (1:3000), polyclonal α-MlaC (Ercan, Low, Liu, & Chng, 2019) (1:500), or α-His antibody (pentahistidine) conjugated to the horseradish peroxidase (HRP) (Qiagen, Hilden, Germany) at 1:3000 dilution for 1 – 3 h at room temperature. Secondary antibody ECL™ anti-mouse IgG-HRP, and anti-rabbit IgG-HRP were used at 1:3000 dilution. Luminata Forte Western HRP Substrate (Merck Millipore) was used to develop the membranes, and chemiluminescence signals were visualized by *G*:Box Chemi-XT4 (Genesys version 1.4.3.0, Syngene).

**AlphaFold2 multimer modelling of MlaA-MlaC complex**

To generate predicted models of the MlaA and MlaC complex, the AlphaFold2 neural-network (Jumper et al., 2021) implemented in the ColabFold pipeline (Mirdita et al., 2022) was used. Using the mature sequences (without signal peptides) of MlaA and MlaC, default options were specified, and multiple sequence alignments produced by MMseqs2 (Mirdita, Steinegger, & Soding, 2019) were used as input for template-free structure prediction by AlphaFold2. Structural relaxation of the final protein geometry was performed using AMBER (Hornak et al., 2006) to obtain five relaxed models of MlaA-MlaC complex. The best scored model was selected and illustrated in cartoon representation using PyMOL (Schrodinger, USA).

**OM permeability studies.** OM sensitivity against SDS/EDTA was judged by colony-forming-unit (CFU) analyses on LB agar plates containing indicated concentrations of SDS/EDTA. Briefly, 5 mL cultures were grown (inoculated with overnight cultures at 1:100 dilution) in LB broth at 37 °C until OD600 reached ~0.4-0.6. Cells were normalized by optical density, first diluted to OD600 = 0.1 (~$10^8$ cells), and then serially diluted (ten-fold) in LB broth using 96-well microtiter plates. 1.5 μL of the diluted cultures were manually spotted onto the plates, dried, and incubated overnight at 37 °C. Plate images were visualized by *G*:Box Chemi-XT4 (Genesys version 1.4.3.0, Syngene).

**SEC-MALS analysis to determine absolute molar masses of OmpC$_3$-MlaA$^{Cys}$-MlaC$^{Cys}$-His complex**

Prior to each SEC–MALS analysis, a preparative SEC was performed for BSA (Sigma–Aldrich) to separate monodisperse monomeric peak and to use as a quality control for the MALS detectors. In each experiment, monomeric BSA was injected before the protein of interest, and the settings (calibration constant for TREOS detector; Wyatt Technology) that gave the well-characterized molar mass of BSA (66.4 kDa) were used for the molar mass calculation of the protein of interest. SEC-purified OmpC$_3$-MlaA$^{Cys}$-MlaC$^{Cys}$-His was concentrated to 3 mg/ml and injected into Superdex 200 Increase 10/300 GL column pre-equilibrated with TBS pH 8.0 and 0.025% DDM. Light scattering and refractive index ($n$) data were collected online using miniDAWN TREOS (Wyatt Technology) and Optilab T-rEX (Wyatt Technology), respectively, and analyzed by ASTRA 6.1.5.22 software (Wyatt Technology). Protein-conjugate analysis available in ASTRA software was applied to calculate non-proteinaceous part of the complex. In this analysis, the refractive index increment $dn/dc$ values (where $c$ is sample concentration) of 0.143 mL/g and 0.185 mL/g were used for DDM and protein complex, respectively (Slotboom, Duurkens, Olieman, & Erkens, 2008). For BSA, UV extinction coefficient of 0.66 ml/(mg.cm) was used. For the OmpC-MlaA-MlaC-His complex, the UV extinction coefficient was calculated to be 1.62 mL/(mg.cm), based on its observed stoichiometric ratio OmpC$_3$-(MlaA$^{Cys}$-MlaC$^{Cys}$-His)$_2$.

**Reconstitution of OmpC$_3$-MlaA$^{Cys}$xMlaC$^{Cys}$-His complex in lipid MSP2N2 nanodiscs**

The reconstitution of purified OmpC$_3$-MlaA$^{Cys}$-MlaC$^{Cys}$-His into nanodiscs was adapted from published protocols (Ritchie et al., 2009). Briefly, 10 mg *E. coli* polar lipid extracts (Avanti Polar Lipids) were dissolved in 1 mL chloroform and dried overnight. Then, 1 mL Tris-buffered saline (TBS) buffer (20 mM Tris HCl pH

8.0, 150 mM NaCl) containing 25 mM sodium cholate (Sigma-Aldrich) was added to the dried lipid film and vortexed and sonicated until a clear solution was obtained. The $OmpC_3$-$MlaA^{Cys}$-$MlaC^{Cys}$-His complex, the membrane scaffold protein MSP2N2-His (Denisov & Sligar, 2016), and lipids were mixed at a molar ratio of 1:2:60 in TBS and incubated, with rocking, for 1 h at 4°C. Bio-beads SM2 resin (Bio-Rad) was subsequently added to the solution (30 mg per 1-mL reconstitution mixture) and incubated with gentle agitation for 1 h at 4°C. Upon removal of Bio-beads, the sample was concentrated and purified by size-exclusion chromatography on a Superose 6 Increase 10/300 GL column (GE Healthcare) on AKTA Pure. 0.5-mL fractions were collected and subjected to reducing (R) and non-reducing (NR) SDS–polyacrylamide gel electrophoresis (SDS-PAGE) and Coomassie blue staining. Fractions containing the nanodisc-embedded complexes were pooled and concentrated on a 100-kDa cut-off ultrafiltration device (Amicon Ultra, Merck Millipore).

**Cryo-EM grid preparation and data acquisition**

For sample preparation, 3.0 μL of the protein sample at a concentration of 12 mg/mL was applied to glow-discharged Quantifoil holey carbon grids (1.2/1.3, 200 mesh). Grids were blotted for 3 s with 100% relative humidity and plunge-frozen in liquid ethane cooled by liquid nitrogen using a Vitrobot System (Gatan). Cryo-EM data were collected at liquid nitrogen temperature on a Titan Krios electron microscope (Thermo Fisher Scientific), equipped with a K3 Summit direct electron detector (Gatan) and GIF Quantum energy filter. All cryo-EM movies were recorded in counting mode with SerialEM4 (Mastronarde, 2005) with a slit width of 20 eV from the energy filter. Movies were acquired at nominal magnifications of 105k, corresponding to a calibrated pixel size of 0.834 Å on the specimen level. The total exposure time of each movie was 6 s, resulting in a total dose of 90 electrons per $Å^2$, fractionated into 50 frames. More details of electron microscopy data collection parameters are listed in **Table S4**.

**Electron microscope image processing**

cryoSPARC 4.0 (Punjani et al., 2017) was used to process the EM data according to the flowchart in **Figure S10**. Dose-fractionated movies were corrected for motion using Patch Motion Correction. To obtain a sum of all frames for each movie, a dose-weighting scheme was applied, and this sum was used for all image-processing steps except for defocus determination. Defocus values of the summed images from all movie frames were calculated using patch CTF estimation without dose weighting. Particle picking was performed using the blob picker followed by the template picker. Two- and three-dimensional (2D and 3D) classifications were carried out using "2D classification", "Ab-initio Reconstruction", and "Heterogeneous Refinement". 3D refinements were conducted using "Homogeneous Refinement" and "Non-Uniform Refinement". Eventually, we obtained three cryo-EM maps representing compositional heterogeneities in $OmpC_3$-MlaA-MlaC complexes, namely $OmpC_3$-(MlaA-MlaC) (**EMD-35253**), $OmpC_3$-$(MlaA\text{-}MlaC)_2$ (**EMD-35252**) and $OmpC_3$-$(MlaA\text{-}MlaC)_3$ (**EMD-35251**). A fourth map for $OmpC_3$-$(MlaA\text{-}MlaC)_{1\text{-}3}$ (**EMD-35250**) was refined, where all particles from the above classes were combined. The overall resolutions for these maps were estimated based on the gold-standard criterion of Fourier shell correlation (FSC) = 0.143, while local resolution was estimated with "Local Resolution Estimation".

**Model building and refinement**

An initial model of $OmpC_3$-*Ec*MlaA was built using the OmpK36-*Kp*MlaA crystal structure (PDB 5NUP) in SWISS-MODEL (Waterhouse et al., 2018). The model was rigid-body fitted into the highest resolution cryo-EM map (**EMD-35250** - 2.93 Å) in Chimera (Pettersen et al., 2004), followed by density-fitting in COOT (Emsley & Cowtan, 2004). Refinement in RealSpace using the program of Phenix (Liebschner et al., 2019) with default parameters yielded the structure for $OmpC_3$-MlaA (**PDB 8I8R**). Next, owing to poor resolution at the MlaC region, a rigid body fitting of MlaC (PDB 5UWA) was first performed with all sidechains removed, albeit in the **EMD-35253** map (3.19 Å resolution, more density at MlaC). After that, an extra density was observed at the distal groove of MlaC from the membrane, which could be clearly traced back to MlaA. A poly-alanine model corresponding to the C-terminal of MlaA was thus built in COOT and refined in RealSpace/Phenix (Liebschner et al., 2019), to yield coordinates for $OmpC_3$-MlaA-MlaC (**PDB 8I8X**). The final coordinates of the asymmetric units were checked using MolProbity (Chen et al., 2010). Maps and structures shown in the figures were generated using UCSF Chimera and COOT. The model building and refinement statistics are shown in **Table S4**.

### Supplementary Tables

**Table S1. Bacterial strains used in this study**

| **Strains** | **Relevant genotypes and characteristics** | **References** |
|---|---|---|
| MC4100 | *F- araD139 Δ(argF-lac) U169 rpsL150 relA1 flbB5301 ptsF25 deoC1 ptsF25 thi* | (Casadaban, 1976) |
| NovaBlue | *endA1 hsdR17 (rK12– mK12+) supE44 thi-1 recA1 gyrA96 relA1 lac F′ proA+ B+ lacIq ZΔM15*::*Tn10* | Novagen |
| BL21(λDE3) | *fhuA2 lon ompT gal (λDE3) dcm* Δ*hsdS* λ*DE3 =* λ *sBamHIo* Δ*EcoRI-B int*::*(lacI*::*PlacUV5*::*T7 gene1) i21* Δ*nin5* | Novagen |
| TKW001 | BL21(λDE3) Δ*ompF*::*kan* | (Yeow et al., 2018) |
| CZS010 | MC4100 Δ*mlaA*::*kan* | (Chong et al., 2015) |
| YJ001 | MC4100 Δ*mlaA*::*FRT* Δ*mlaC*::*kan* | This study |
| WY001 | MC4100 Δ*mlaA::FRT* Δ*mlaD*::*kan* | (Ercan et al., 2019) |
| CZS015 | MC4100 Δ*ompC*::*kan* | (Chong et al., 2015) |
| YJ002 | MC4100 Δ*ompC::FRT* Δ*mlaC*::*kan* | This study |
| CZS608 | MC4100 Δ*ompC*::*ompC*$_{R92A}$; chromosomal *ompC* mutation introduced via a positive-negative selection | (Khetrapal et al., 2015; Yeow et al., 2018) |
| RS112 | MC4100 Δ*tolB*::*kan* | (Shrivastava, Jiang, & Chng, 2017) |
| RS123 | MC4100 Δ*pal*::*kan* | (Shrivastava et al., 2017) |

**Table S2. Plasmids used in this study**

| Plasmids | Relevant genotypes and characteristics | References |
|---|---|---|
| pET22/42 | pT7(*lac*$_O$) inducible expression vector, contains multiple cloning site of pET42a(+) in pET22b(+) backbone; Amp$^R$ | (Wu et al., 2006) |
| pET23/42 | pT7 inducible expression vector, contains multiple cloning site of pET42a(+) in pET23a(+) backbone; Amp$^R$ | (Wu et al., 2006) |
| pSup-BpaRS-6TRN | Encodes an orthogonal tRNA and aminoacyl-tRNA synthetase permitting ribosomal incorporation of *p*Bpa at TAG stop codons (pSup) | (Ryu & Schultz, 2006) |
| pCDFDuet-1 | pT7 inducible expression vector; Spec$^R$ | Novagen |
| pDSW206 | Promoter down mutations in -35 and -10 of pTrc99a; Amp$^R$ | (Weiss, Chen, Ghigo, Boyd, & Beckwith, 1999) |
| pET23/42*mlaA-His* | Encodes full length MlaA with C-terminal His8 tag; Amp$^R$ | (Chong et al., 2015) |
| pET23/42*mlaA*$_{3G3P}$-*His* | Encodes full length MlaA$_{3G3P}$ with C-terminal His8 tag; Amp$^R$ | (Yeow et al., 2018) |
| pET23/42*mlaA**-*His* | Encodes full length MlaA* with C-terminal His8 tag; Amp$^R$ | (Yeow et al., 2018) |
| pCDF*mlaA-His* | Encodes full length MlaA with C-terminal His8 tag; Spec$^R$ | (Yeow et al., 2018) |
| pCDF*mlaC*$^{V171C}$-*His-mlaA*$^{Q205C}$ | Encodes full length MlaC$^{V171C}$ with C-terminal His6 tag and full length MlaA$^{Q205C}$; Spec$^R$ | This study |
| pET23/42*mlaC-mlaB* | Encodes full length MlaC and MlaB; Amp$^R$ | Feifan Zhu |
| pET22/42*mlaC-His* | Encodes full length MlaC with C-terminal His8 tag; Amp$^R$ | (Ercan et al., 2019) |
| pDSW206*ompC* | Encodes full length OmpC; Amp$^R$ | (Yeow et al., 2018) |
| pDSW206*ompF* | Encodes full length OmpF; Amp$^R$ | (Yeow et al., 2018) |
| pDSW206*ompC-His* | Encodes full length OmpC with C-terminal His8 tag; Amp$^R$ | (Yeow et al., 2018) |
| pTrc99a-*pldA* | Encodes full length PldA; Amp$^R$ | (Chong et al., 2015) |

**Table S3. Primers used in this study.**

| Primers | Sequence (5' to 3')* |
|---|---|
| mlaA_R28B FP | GATCAGCAAGGG tag TCTGACCCGTTAGAAGGGTTC |
| mlaA_R28B RP | CTAACGGGTCAGA cta CCCTTGCTGATCTGTACCGG |
| mlaA_N41B FP | GCACCATGTAC tag TTCAACTTCAATGTATTAG |
| mlaA_N41B RP | TTGAAGTTGAA cta GTACATGGTGCGGTTGAAC |
| mlaA_N45B FP | ACTTCAACTTC tag GTATTAGACCCGTATATTG |
| mlaA_N45B RP | GGGTCTAATAC cta GAAGTTGAAGTTGTACATG |
| mlaA_P49B FP | AATGTATTAGAC tag TATATTGTTCGACCGGTC |
| mlaA_P49B RP | CGAACAATATA cta GTCTAATACATTGAAGTTG |
| mlaA_Y50B FP | TATTAGACCCG tag ATTGTTCGACCGGTCGCTG |
| mlaA_Y50B RP | GGTCGAACAAT cta CGGGTCTAATACATTGAAG |
| mlaA_P54B FP | TATATTGTTCGA tag GTCGCTGTCGCCTGGCG |
| mlaA_P54B RP | GGCGACAGCGAC cta TCGAACAATATACGGGTC |
| mlaA_A58B FP | CCGGTCGCTGTC tag TGGCGTGATTATGTTCCG |
| mlaA_A58B RP | ATAATCACGCCA cta GACAGCGACCGGTCGAAC |
| mlaA_D61B FP | GTCGCCTGGCGT tag TATGTTCCGCAACCGGCG |
| mlaA_D61B RP | TTGCGGAACATA cta ACGCCAGGCGACAGCGAC |
| mlaA_A194B FP | GATCGAAACCCGC tag CAGCTGCTGGATTCCGATG |
| mlaA_A194B RP | ATCCAGCAGCTG cta GCGGGTTTCGATCCCTTCAAG |
| mlaA_Q195B FP | GAAACCCGCGCT tag CTGCTGGATTCCGATGG |
| mlaA_Q195B RP | GAATCCAGCAG cta AGCGCGGGTTTCGATCCC |
| mlaA_L196B FP | CCCGCGCTCAG tag CTGGATTCCGATGGTCTGC |
| mlaA_L196B RP | CATCGGAATCCAG cta CTGAGCGCGGGTTTCGATC |
| mlaA_L197B FP | CGCGCTCAGCTG tag GATTCCGATGGTCTGCTG |
| mlaA_L197B RP | CCATCGGAATC cta CAGCTGAGCGCGGGTTTCG |
| mlaA_D198B FP | GCTCAGCTGCTG tag TCCGATGGTCTGCTGCGTC |
| mlaA_D198B RP | CAGACCATCGGA cta CAGCAGCTGAGCGCGGG |
| mlaA_S199B FP | CGCTCAGCTGCTGGAT tag GATGGTCTGC |
| mlaA_S199B RP | CTGACGCAGCAGACCATC cta ATCCAGCAGC |
| mlaA_D200B FP | CTCAGCTGCTGGATTCC tag GGTCTGCTGCG |
| mlaA_D200B RP | CGACTGACGCAGCAGACC cta GGAATCCAGCAG |
| mlaA_G201B FP | CTGGATTCCGAT tag CTGCTGCGTCAGTCGTCCG |
| mlaA_G201B RP | CTGACGCAGCAG cta ATCGGAATCCAGCAGCTG |
| mlaA_L202B FP | GATTCCGATGGT tag CTGCGTCAGTCGTCCG |
| mlaA_L202B RP | GACTGACGCAG cta ACCATCGGAATCCAGCAGC |
| mlaA_L203B FP | CCGATGGTCTG tag CGTCAGTCGTCCGATCC |
| mlaA_L203B RP | GACGACTGACG cta CAGACCATCGGAATCCAGC |
| mlaA_R204B FP | GATTCCGATGGTCTGCTG tag CAGTCGTCCGATCC |
| mlaA_R204B RP | AATATAAGGATCGGACGACTG cta CAGCAGACCATC |
| mlaA_Q205B FP | CCGATGGTCTGCTGCGT tag TCGTCCGATCC |
| mlaA_Q205B RP | CATAATATAAGGATCGGACGA cta ACGCAGCAGAC |
| mlaA_D208B FP | CGTCAGTCGTCC tag CCTTATATTATGGTGCGC |
| mlaA_D208B RP | CATAATATAAGG cta GGACGACTGACGCAGCAG |
| mlaA_M212B FP | GATCCTTATATT tag GTGCGCGAAGCGTACTTC |
| mlaA_M212B RP | CGCTTCGCGCAC cta AATATAAGGATCGGACGAC |
| mlaA_E215B FP | TTATGGTGCGC tag GCGTACTTCCAGCGTCATG |
| mlaA_E215B RP | CTGGAAGTACGC cta GCGCACCATAATATAAGG |
| mlaA_A216B FP | ATGGTGCGCGAA tag TACTTCCAGCGTCATGAT |
| mlaA_A216B RP | ACGCTGGAAGTA cta TTCGCGCACCATAATATA |

| | |
|---|---|
| mlaA_Q219B FP | CGAAGCGTACTTC tag CGTCATGATTTCATCGC |
| mlaA_Q219B RP | AAATCATGACG cta GAAGTACGCTTCGCGCACC |
| mlaA_D222B FP | TTCCAGCGTCAT tag TTCATCGCTAATGGCGGC |
| mlaA_D222B RP | ATTAGCGATGAA cta ATGACGCTGGAAGTACGC |
| mlaA_F223B FP | AGCGTCATGAT tag ATCGCTAATGGCGGCGAAC |
| mlaA_F223B RP | CCATTAGCGAT cta ATCATGACGCTGGAAGTAC |
| mlaA_Q195C FP | GAAACCCGCGCT tgc CTGCTGGATTCCGATGG |
| mlaA_Q195C RP | GAATCCAGCAG gca AGCGCGGGTTTCGATCCC |
| mlaA_Q205C FP | CCGATGGTCTGCTGCGT tgc TCGTCCGATCC |
| mlaA_Q205C RP | CATAATATAAGGATCGGACGA gca ACGCAGCAGAC |
| mlaA_M212C FP | GATCCTTATATT tgc GTGCGCGAAGCGTACTTC |
| mlaA_M212C RP | CGCTTCGCGCAC gca AATATAAGGATCGGACGA |
| mlaA_F223C FP | AGCGTCATGAT tgc ATCGCTAATGGCGGCGAAC |
| mlaA_F223C RP | CCATTAGCGAT gca ATCATGACGCTGGAAGTAC |
| mlaC_V171C_FP | CATGATTGCTGAAGGC tgc AGTATGATCACCAC |
| mlaC_V171C_RP | ATCATACT gca GCCTTCAGCAATCATGTCGTAA |
| mlaC_K128C_FP | CCGCTGGGCGAT tgc ACCATTGTGCCTATTCGC |
| mlaC_K128C_RP | AGGCACAATGGT gca ATCGCCCAGCGGCTGTTC |
| mlaC_P132C_FP | AAACCATTGTG tgc ATTCGCGTTACCATTATTG |
| mlaC_P132C_RP | TGGTAACGCGAAT gca CACAATGGTTTTATCGC |
| mlaC_Q151C_FP | CGTCTGGACTTC tgc TGGCGTAAAAACTCCCAG |
| mlaC_Q151C_RP | GTTTTTACGCCA gca GAAGTCCAGACGCACCGG |
| mlaC_S172C_FP | TTGCTGAAGGCGTC tgc ATGATCACCACCAAAC |
| mlaC_S172C_RP | GTGGTGATCAT gca GACGCCTTCAGCAATCATG |
| mlaC_T175C_FP | GTCAGTATGATC tgc ACCAAACAAAACGAGTGG |
| mlaC_T175C_RP | GTTTTGTTTGGT gca GATCATACTGACGCCTTC |
| mlaA_CTD4 FP | GGCGAACTCAAA taa CAGGAAAATCCGAACGC |
| mlaA_CTD4 RP | GGATTTTCCTG tta TTTGAGTTCGCCGCCATT |
| mlaA_CTD5 FP | CAGGAAAATCCG taa GCACAAGCGATTCAGGATG |
| mlaA_CTD5 RP | GAATCGCTTGTGC tta CGGATTTTCCTGCGGTTTG |
| mlaA_CTD6 FP | CAAGCGATTCAG taa GATTTAAAAGATATTG |
| mlaA_CTD6 RP | CTTTTAAATC tta CTGAATCGCTTGTGCGTTC |
| mlaA_2DD2R FP | CAAGCGATTCAG cgc cgc TTAAAAGATATTGATTC |
| mlaA_2DD2R RP | ATATCTTTTAA gcg gcg CTGAATCGCTTGTGCGTTC |
| mlaA_2DE2R FP | GATTTAAAA cgc ATTGATTCT cgc CTCGAGCACCACC |
| mlaA_2DE2R RP | GCTCGAG gcg AGAATCAAT gcg TTTTAAATCATCCTG |
| mlaC0 FP | CGTACAGGTG gac TACGCCGGTGCGCTG |
| mlaC0 RP | ACCGGCGTA gtc CACCTGTACGTATGGC |
| mlaC1 FP | GTATTAC gac AGTGCGACCCCTGCTCAA gac GAAGCCTACT |
| mlaC1 RP | GCTTC gtc TTGAGCAGGGGTCGCACT gtc GTAATACTGGCC |
| mlaC2 FP | CCGCTTTC gac GAGTACCTG gac CAGGCTTACGGTCAG |
| mlaC2 RP | CGTAAGCCTG gtc CAGGTACTC gtc GAAAGCGGCAAAG |
| mlaC3 FP | ATTGACCCGAATGGC gac CCGCCGGTG gac CTGGACTTCC |
| mlaC3 RP | CCACTGGAAGTCCAG gtc CACCGGCGG gtc GCCATTCGGG |
| mlaC4 FP | CTTCCAGTGG gac gac AACTCCCAGACGGGCAAT |
| mlaC4 RP | CGTCTGGGAGTT gtc gtc CCACTGGAAGTCCAG |
| mlaC5 FP | GGAACGCTGCTG gac ACC gac GGTATCGACGGCCTG |
| mlaC5 RP | CGTCGATACC gtc GGT gtc CAGCAGCGTTCCCCACTC |
| mlaC7 FP | AACTG gac TCGATTTCTCAACAG gac ATCACTCTGGAAG |
| mlaC7 RP | GAGTGAT gtc CTGTTGAGAAATCGA gtc CAGTTGCGCAG |
| mlaA_M165C FP | GACGGTGGTGAT tgc GCGGATGGTTTTTACCCGG |

| | |
|---|---|
| mlaA_M165C RP | AAAACCATCCGC gca ATCACCACCGTCATCACGC |
| mlaA_A166C FP | GGTGGTGATATG tgc GATGGTTTTTACCCGGTTC |
| mlaA_A166C RP | GTAAAAACCATC gca CATATCACCACCGTCATC |
| mlaA_D167C FP | GACGGTGGTGATATGGCG tgc GGTTTTTACCCG |
| mlaA_D167C RP | AAGAACCGGGTAAAAACC gca CGCCATATCACC |
| mlaA_Y170C FP | GCGGATGGTTTT tgc CCGGTTCTTTCCTGGCTG |
| mlaA_Y170C RP | GGAAAGAACCGG gca AAAACCATCCGCCATATC |
| mlaA_V172C FP | GGTTTTTACCCG tgc CTTTCCTGGCTGACCTGG |
| mlaA_V172C RP | CAGCCAGGAAAG gca CGGGTAAAAACCATCCGC |
| mlaA_L173C FP | GATGGTTTTTACCCGGTT tgc TCCTGGCTGACC |
| mlaA_L173C RP | CGGCCAGGTCAGCCAGGA gca AACCGGGTAAAA |
| mlaA_S174C FP | TACCCGGTTCTT tgc TGGCTGACCTGGCCGATG |
| mlaA_S174C RP | CCAGGTCAGCCA gca AAGAACCGGGTAAAAACC |
| mlaA_W175C FP | TTTTACCCGGTTCTTTCC tgc CTGACCTGGCCG |
| mlaA_W175C RP | CAGACATCGGCCAGGTCAG gca GGAAAGAACCGG |
| mlaA_L176C FP | CTTTCCTGG tgc ACCTGGCCGATGTCTGTG |
| mlaA_L176C RP | CATCGGCCAGGT gca CCAGGAAAGAACCGGG |
| mlaA_T177C FP | CTTTCCTGGCTG tgc TGGCCGATGTCTGTGGG |
| mlaA_T177C RP | GACATCGGCCA gca CAGCCAGGAAAGAACCGG |
| mlaA_W178C FP | GTTCTTTCCTGGCTGACC tgc CCGATGTCTGTG |
| mlaA_W178C RP | ATTTACCCACAGACATCGG gca GGTCAGCCAGG |
| mlaA_P179C FP | GGCTGACCTGG tgc ATGTCTGTGGGTAAATGG |
| mlaA_P179C RP | CCCACAGACAT gca CCAGGTCAGCCAGGAAAG |
| mlaA_M180C FP | CTGACCTGGCCG tgc TCTGTGGGTAAATGGACG |
| mlaA_M180C RP | TTTACCCACAGA gca CGGCCAGGTCAGCCAGG |
| mlaA_S181C FP | CTGGCCGATG tgc GTGGGTAAATGGACGCTTG |
| mlaA_S181C RP | CATTTACCCAC gca CATCGGCCAGGTCAGCCAG |
| mlaA_W185C FP | CTGTGGGTAAA tgc ACGCTTGAAGGGATCGAAAC |
| mlaA_W185C RP | CCCTTCAAGCGT gca TTTACCCACAGACATCG |
| mlaA_L187C FP | GGTAAATGGACG tgc GAAGGGATCGAAACCCGCG |
| mlaA_L187C RP | GTTTCGATCCCTTC gca CGTCCATTTACCCACAGAC |
| mlaA_E188C FP | AAATGGACGCTT tgc GGGATCGAAACCCGCGC |
| mlaA_E188C RP | GTTTCGATCCC gca AAGCGTCCATTTACCCAC |
| mlaA_R193C FP | CTTGAAGGGATCGAAACC tgc GCTCAGCTGCTG |
| mlaA_R193C RP | GGAATCCAGCAGCTGAGC gca GGTTTCGATCCC |

**Table S4. Summary of cryo-EM data collection and model refinement.**

| **Dataset Collection** | | | | |
|---|---|---|---|---|
| Voltage (kV) | 300 | | | |
| Magnification | 105,000x | | | |
| Calibrated pixel size (Å) | 0.834 | | | |
| Defocus range (µm) | -0.8 to -2.0 | | | |
| Total electron dose (e-/Å$^2$) | 90 | | | |
| Number of images | 6018 | | | |
| Number of frames/image | 50 | | | |
| **Maps** | **OmpC$_3$-(MlaA-MlaC)$_{1-3}$** | **OmpC$_3$-(MlaA-MlaC)$_3$** | **OmpC$_3$-(MlaA-MlaC)$_2$** | **OmpC$_3$-(MlaA-MlaC)** |
| EMDB ID | 35250 | 35251 | 35252 | 35253 |
| # Particles used | 205,487 | 38,528 | 90,496 | 76,463 |
| Resolution (Å) (0.143 threshold) | 2.93 | 3.42 | 3.19 | 3.25 |
| B-factor sharpening | 88.0 | 50.2 | 73.8 | 69.1 |
| **Refinement coordinates** | **OmpC-MlaA** | | | **OmpC-MlaA-MlaC** |
| PDB ID | 8I8R | | | 8I8X |
| **Composition** | | | | |
| # Chains | 7 | | | 8 |
| # Residues | 1,232 | | | 1,450 |
| # Non-H Atoms | 10,151 | | | 11,226 |
| # Ligands | 3 | | | 3 |
| **Bonds (RMSD)** | | | | |
| Length (Å) | 0.002 | | | 0.007 |
| Angles (°) | 0.622 | | | 1.174 |
| MolProbity score | 1.73 | | | 1.92 |
| Clash score | 4.14 | | | 4.40 |
| **Ramachandran plot (%)** | | | | |
| Outliers | 0.16 | | | 0.28 |
| Allowed | 7.11 | | | 7.71 |
| Favored | 92.73 | | | 92.01 |
| **Rama-Z (Z-score, RMSD)** | | | | |
| Whole (#) | -2.70 (0.22), 1224 | | | -3.26 (0.20), 1440 |
| Helix (#) | -3.30 (0.27), 151 | | | -3.88 (0.15), 236 |
| Sheet (#) | -0.34 (0.24), 495 | | | -0.50 (0.23), 523 |
| Loop (#) | -2.73 (0.21), 578 | | | -2.90 (0.20), 681 |
| Rotamer outliers (%) | 1.82 | | | 0.71 |
| Cβ deviations (%) | 0.00 | | | 0.08 |
| **Peptide plane (%)** | | | | |
| Cis proline/general | 0.0/0.0 | | | 0.0/0.0 |
| Twisted proline/general | 0.0/0.0 | | | 0.0/0.0 |
| CaBLAM outliers (%) | 4.52 | | | 5.17 |
